# Seed banks alter the molecular evolutionary dynamics of *Bacillus subtilis*

**DOI:** 10.1101/2021.10.05.463161

**Authors:** William R. Shoemaker, Evgeniya Polezhaeva, Kenzie B. Givens, Jay T. Lennon

**Affiliations:** Department of Biology, Indiana University, Bloomington, IN, 47405, USA; Department of Ecology and Evolutionary Biology, UCLA, Los Angeles, CA, 90095, USA; Luddy School of Informatics, Computing, and Engineering, Indiana University, Bloomington IN, 47408, USA

## Abstract

Fluctuations in the availability of resources constrains the growth and reproduction of individuals, which subsequently effects the evolution of their respective populations. Many organisms contend with such fluctuations by entering a reversible state of reduced metabolic activity, a phenomenon known as dormancy. This pool of dormant individuals (i.e., a seed bank) does not reproduce and is expected to act as an evolutionary buffer, though it is difficult to observe this effect directly over an extended evolutionary timescale. Through genetic manipulation, we analyze the molecular evolutionary dynamics of *Bacillus subtilis* populations in the presence and absence of a seed bank over 700 days. The ability of these bacteria to enter a dormant state increased the accumulation of genetic diversity over time and altered the trajectory of mutations, findings that were recapitulated using simulations based on a mathematical model of evolutionary dynamics. While the ability to form a seed bank did not alter the degree of negative selection, we found that it consistently altered the direction of molecular evolution across genes. Together, these results show that the ability to form a seed bank can affect the direction and rate of molecular evolution over an extended evolutionary timescale.

## Introduction

Nature is rarely static. Temporal fluctuations in abiotic and biotic environmental factors often reduce the rate that an organism can grow and reproduce. To contend with such fluctuations, many species enter a reversible state of reduced metabolic activity, an adaptation known as dormancy [1]. In this state, individuals can endure environmental stressors until they subside, a temporary cessation of short-term reproductive efforts to increase long-term reproductive gains. This evolutionary trade-off, and the life-history strategies through which it is implemented, have received substantial attention by means of theoretical [2–7] and empirical investigations [8–10], spurred in-part by the observation that dormancy has independently evolved multiple times across the tree of life [11–13].

While the trade-off aspect of dormancy has been of considerable historical interest, life-history traits do not operate in a population genetic vacuum. The fitness benefit of a life-history trait is often a consequence of its effect on a given birth-death process [14], population dynamics that sequentially alter the dynamics and fates of genetic variants. The ability to enter a dormant state is no exception. The accumulation of dormant individuals within a system can result in the formation of seed banks [11], demographic structures that can reshape the molecular evolutionary dynamics of a population.

Seed banks primarily alter the molecular evolutionary dynamics of a population through two means. First, the ability to enter a dormant state dampens the accumulation of *de novo* genetic diversity as well as its fluctuations over its sojourn time, as dormant individuals do not reproduce and the vast majority of mutations are typically acquired during the process of genome replication [15, 16]. Second, seed banks can act as reservoirs of genetic and phenotypic diversity. These reservoirs reduce the efficiency of natural selection [17–19], dampen the loss of genetic diversity due to random genetic drift [19–24], and permit the retention of deleterious variants [25, 26]. Together, the presence of a seed bank reduces the rate of molecular evolution while increasing the maximum amount of genetic diversity that can be retained. Furthermore, because the ability to form a seed bank is the result of a life-history strategy maintained by natural selection, it is possible that the formation of seed banks restricts the targets of molecular evolution, constraining the direction of evolution as well as its rate. While substantial progress has been made towards developing mathematical models that describe these patterns within the discipline of theoretical population genetics [18–22], there remains a comparative lack of experimental tests of central predictions.

The role of seed banks as an evolutionary buffer means that time is an essential factor when considering an appropriate empirical system. Namely, if the per-generation rate of change of genetic diversity is reduced by a given amount it is often necessary to observe an additional proportionate number of generations. This constraint makes it challenging to directly observe seed bank dynamics over extended evolutionary timescales. Given their short generation times, large population sizes, propensity for rapid adaptation, and the prevalence of dormancy among their lineages [1], microorganisms are an ideal group of organisms to characterize the extent that dormancy alters molecular evolutionary dynamics. In addition, certain lineages of microorganisms have evolved the ability to form complex protective structures (i.e., endospores) that allow them to survive and form long-lasting seed banks [27, 28]. While these structures are not the only means by which microorganisms can enter a dormant state [29], their existence provides a means through which the formation of seed banks can be genetically manipulated [30]. Furthermore, while evolution experiments have been performed using seed bank forming microorganisms [31], questions pertaining to dormancy have primarily been restricted to examining the phenotypic decay of endospore formation via the acquisition of *de novo* mutations under relaxed selection [32–34], whereas the effect of endospore formation on the molecular evolutionary dynamics of microorganisms has remained relatively unexplored. In this study, we examined the molecular evolutionary dynamics of *Bacillus subtilis* populations that differ in their ability to form protective endospores, a non-reproductive structure that is the primary mechanism through which this species enters a dormant state to form a seed bank. To manipulate seed bank formation we replenished the resources of dormancy-capable and incapable populations via serial dilution every 1, 10 and 100 days, a set of transfer regimes that allowed us to examine the effect of dormancy across a range of energy availabilities. Replicate populations were maintained for over 700 days, generating a molecular fossil record which was reconstructed to determine how the presence of a seed bank altered the trajectories of *de novo* mutations. We then recapitulated the dynamics we observed using simulations based on a stochastic model of molecular evolution in dormancy-capable populations. Finally, we identified the sets of genes that were enriched for mutations within each transfer-regime for dormancy-capable and incapable populations, allowing us to quantify parallel evolution among replicate populations as well as the degree of divergent evolution between dormancy-capable and incapable populations.

## Materials and Methods

### Mutant construction

To manipulate endospore formation we deleted *spo0A*, the master regulatory gene for sporulation pathways in *B. subtilis*. Gene deletion was performed using Gibson assembly of PCR amplified dsDNA fragments upstream and downstream of *spo0A* (Table S1). Purified ligated plasmid was transformed into *E. coli* DH5*α* and plasmid DNA was purified from cultured positive transformants. Purified plasmid product was transformed into *E. Coli* TG1, positive transformants were selected, and plasmid DNA was purified before a single *B. subtilis* NCIB 3610 colony was grown in medium containing a purified plasmid aliquot and identified using antibiotic plating. Transformation was confirmed via Sanger sequencing of PCR products and loss of antibiotic resistance was confirmed via antibiotic plating (see Supplemental Methods for additional detail).

### Evolution experiment

#### Fitness assay

We performed fitness assays to determine the degree that the fitness effect of endospore formation varied across transfer-regimes. At mid-exponential phase, equal ratio aliquots of *B. subtilis* WT and *B. subtilis* Δ*spo0A* cultures were transferred to three replicate flasks with fresh medium. Aliquots were frequently sampled from each flask in a sterile biosafety cabinet over 100 days. Strains were distinguishable by the morphology of their Colony Forming Units (CFUs) on agar plates, which were used to obtain estimates of the size of the population (*N*) in a flask via serial dilution. To make sure the CFU morphology of a given strain was stable, we maintained five replicate populations of WT and Δ*spo0A* strains in separate flasks for 100 days and plated each population every 10 days. We found zero evidence that WT CFUs converted to Δ*spo0A* morphology or that Δ*spo0A* CFUs converted to WT morphology, allowing us to use colony morphology to distinguish between strains for a 100 day fitness assay. The relative log fitness of Δ*spo0A* after *t* days was defined as

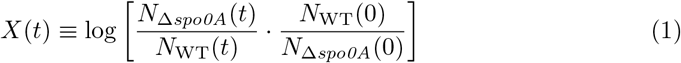

Because we were measuring populations that exited their exponential phase of growth, we do not know how many generations accrued over the course of the fitness assay. Therefore, we chose to examine *X* at each given time point rather than attempt the typical fitness per-generation estimate 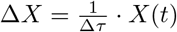.

#### Transfer protocol, sequencing, and variant calling

To evaluate how environmental conditions interacted with the ability to form seed banks, we manipulated energy limitation by extending the time between transfers for microbial populations. We performed our energy-limited evolution experiments using *Bacillus subtilis* NCIB 3610 Δ*spo0A* (ASM205596v1) using a previously described methodology [28]. To briefly summarize, a single colony was isolated from each strain and grown overnight in 10 mL of PYE medium with 0.2% glucose and 0.1% casamino acids in a 50 mL Erlenmeyer flask and split into replicate populations. Five replicates were transferred as 1 mL aliquots into 9 mL of medium every 1, 10, or 100 days for 700 days at 25°C and 250 RPM. All replicate populations were cryopreserved every 100 days by mixing biomass with 20% glycerol solution. These samples were flash-frozen in liquid nitrogen and stored at −80°C. Biomass for DNA extraction was collected from all replicate populations every 100 days (Fig. S1). Populations were regularly plated to test for contamination. An identical experiment was concurrently run with the WT strain, which was previously described [35]. To estimate the number of endospore CFUs, we performed heat treatment at 65°C on aliquots of cultures to kill vegetative cells. CFU counts of vegetative cells were then back-calculated using total CFU counts from non-heat treated culture (i.e., CFU_veg_ = CFU_all_ – CFU_spore_).

In our chosen 1:10 dilution ratio transfer regime under the assumption that the decline in population size was negligible between transfers, we can calculate the number of generations that occurred between a given transfer as Δ*τ* = log_2_ (Dilution factor) = log_2_(10) ≈ 3.3. However, a substantial number of cells died in the 10 and 100-day transfer regimes between transfer events for both WT and Δ*spo0A* strains, meaning that the final size of a population (*N_f_*) relative to its initial size immediately after a transfer (*N_i_*) was not equal to a dilution factor of 10. Thus, estimates of Δ*τ* obtained from the dilution factor will underestimate the true number of generations due to exponential growth per-transfer. To account for this discrepancy, we calculated the number of generations per-transfer using the equivalent notation Δ*τ* = log_2_(*N_f_*/*N_i_*), where *N_f_*/*N_i_* > Dilution factor > 10 in the 10 and 100-day transfer regimes (1). Estimates of N were obtained via CFU counts. These estimates were used for calculating all quantities that contain “generations” as a unit (Eq. 3, 4b).

While this estimate is more accurate than the one based on dilution factors, it does not account for the number of generations that accrued between transfers after a population exited its exponential phase of growth. This unknown number of generations likely occurs as a result of living cells using dead cells as a resource for reproduction. While there is no reliable approach to calculate the number of generations once a population has exited exponential growth, known as “cryptic growth” [35–37], evidence suggests that this number is likely low [28, 38] and can be safely neglected for the purpose of this study.

DNA extraction, library preparation, and pooled population sequencing was performed on all Δ*spo0a* and WT timepoints for all replicate populations as previously described [35]. The first 20 bp of all reads were trimmed and all read pairs where at least one pair had a mean Phred quality score (= −10log_10_*P*; *P* = probability of an incorrect base call) less than 20 were removed via `cutadept v1.9.1` [39]. Candidate variants were identified using a previously published approach [40] that relied on alignments generated from `breseq v0.32.0` [41], which was modified as previously described [35].

#### Mutation trajectory analyses

We estimated the frequency of the *m*th mutation candidate in the *p*th population at the *t*th timepoint using the naive estimator 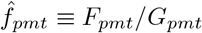, where *F_pmt_* and *G_pmt_* are the total number of reads containing the alternate allele and the total depth of coverage, respectively. We examined the accumulation of mutations by time *t* in the *p*th population as the sum of derived allele frequencies

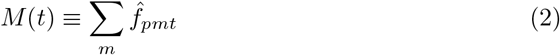

Given that the log of *M*(*t*) over time often appeared to saturate in this study as well as in previous studies [35, 40], we modelled the relationship between *M*(*t*) and *t* using the following equation.

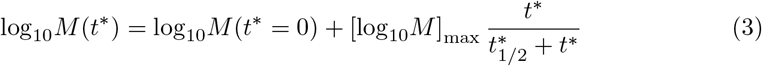

Where [log_10_*M*]_max_ is the maximum value of log_10_*M*(*t*\*) and 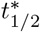 is the value of *t*\* where log_10_*M*(*t*\*) is half of [log_10_*M*]_max_. The variable *t*\* represents the shift in time so that Eq.3 reduces to the intercept parameter (log_10_*M*(0)) at the first temporal sample, in this case, *t*\* = *t* – 100 days. We then multiplied 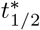 by the estimated minimum number of per-day generations, the product of which we define as *τ*_1/2_. While this model is phenomenological in that we do not posit a microscopic mechanism, much like the Michaelis–Menten kinetic model from which it is derived [42], it effectively captures the hyperbolic pattern. Numerical optimization was performed over 54 initial conditions using the BFGS algorithm in Python using `statsmodels` [43]. Estimates of the probability of extinction of alleles that were detected Pr[Extinct|Detected] for a given strain-transfer regime combination were calculated as the number of mutations where an extinct state was inferred by the end of the experiment divided by the total number of detected mutations.

We examined three different measures to determine how the ability to enter a dormant state affected molecular evolutionary dynamics, defined as

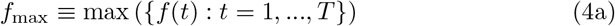

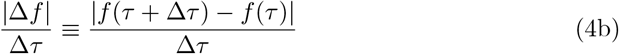

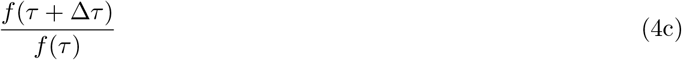

First, *f_max_* is the maximum estimated frequency of a given mutation over *T* observations. The majority of replicate populations had *T* = 7 observations. Because certain 100-day transfer regime replicate populations had *T* = 6 observations, we only used these six observations for estimates of *f_max_* for all replicate populations. Second, |Δ*f*| /Δ*τ* is the magnitude of change in *f* between two observations. Finally, *f* (*τ* + Δ*τ*)/*f* (*τ*) is the direction of change for *f* between two observations. Given the absence of information about physical linkage between mutations in pooled data, we calculated a measure of statistical dependence based on a previously described implementation [44]. As a measure of the rate of recombination, we calculated the Pearson correlation coefficient between all pairs of mutations that were simultaneously segregating (i.e., 0 < *f* < 1) for at least three timepoints for each replicate population.

We compared the empirical cumulative distribution functions of WT and Δ*spo0A* for all measures for each transfer regime using the Kolmogorov–Smirnov (KS) test. To identify fixed mutations, we used a previously published hidden Markov model [40] to infer whether a given mutation eventually became fixed within a replicate population over the course of the experiment. The ratio of nonsynonymous to synonymous polymorphic mutations *pN/pS* was calculated in each population, where the total number of observed mutations of each class was weighted by the relative frequency of nonsynonymous and synonymous sites in all genes in the genome.

### Parallelism at the gene level

We identified potential targets of selection by examining the distribution of nonsynonymous mutations across genes using a previously published approach [40]. To briefly summarize, gene-level parallelism was assessed by calculating the excess number of nonsynonymous mutations acquired at a given gene, relative to the expectation if mutation accumulation was primarily driven by gene size. This quantity, known as *multiplicity*, is defined as

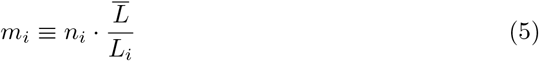

where *n_i_* is the number of mutations observed in the *i*th gene across all five replicate populations for a given strain-transfer regime combination. The term *L_i_* is the effective number of nonsynonymous bases in the gene and 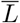 is the mean number of nonsynonymous bases of all genes. Under this definition, the null hypothesis is that all genes have the same multiplicity 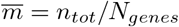. Using the observed and expected values, we quantified the net increase of the log-likelihood of the alternative hypothesis relative to the null

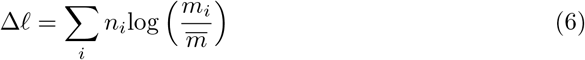

where significance was assessed using permutation. The null hypothesis that genes acquired a random number of nonsynonymous mutations can be captured through the Poisson distribution [40]. Using Eq. 5, we defined the single parameter of the Poisson null as 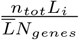. To identify specific genes that are enriched for mutations, we calculated the *P*-value of each gene using the Poisson distribution as

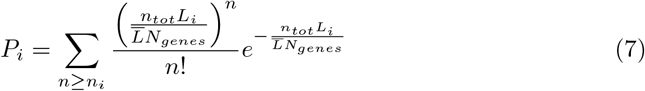

where False Discovery Rate (FDR) correction was performed by defining a critical *P*-value (*P*\*) based on the survival curve of a null Poisson distribution. To increase statistical power we only examined a gene within a given strain-transfer regime combination if it accrued at least three mutations across all replicate populations *n_i_* ≥ 3.

We then defined the set of significant genes for each strain-transfer combination for *α* = 0.05 as:

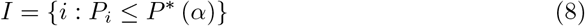

To determine whether genes associated with the *spo0A* regulon were enriched for nonsynonymous mutations, we used an approach similar to Eq. 7. The question of whether a set of genes acquired more mutations than expected by chance can be modeled as a binomial process, where the probability that a given region acquires a mutation is equal its fraction of the genome (*l* = *L*_region_/*L*_genome_). For *spo0A*, *l_spo0a_* ≈ 0.03. Using this model, we define the following P-value.

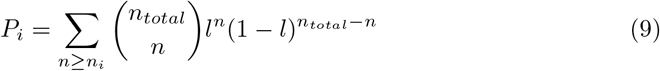

### (Con/di)vergence at the gene level

We tested for convergent vs. divergent evolution by identifying the set of genes that were enriched within a given combination of two strain-transfer regimes (*I*_1_ ∩ *I*_2_). The null distribution for the size of *I*_1_ ∩ *I*_2_ can be modelled as the process of sampling a given number of genes out of *N_genes_* in the genome without replacement two times, a sampling process that is captured by the hypergeometric distribution [45]. By extending this distribution to the multivariate case [35], we obtained a null probability distribution for the number of genes that are enriched in a given pair of strain-transfer regimes

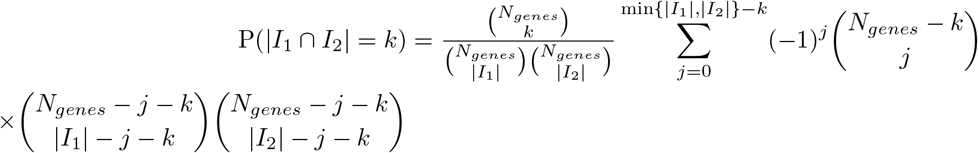

Using this distribution, we determined whether the number of intersecting enriched genes is greater than (convergence) or less than (divergence) our null expectation.

To examine convergent/divergent evolution among enriched genes, we calculated the vector of relative multiplicities 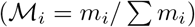 and compared the mean absolute difference between *I* genes for a given pair of transfer regimes or strains as

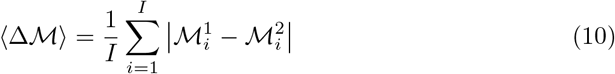

Null distributions of 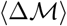 were generated by constructing a gene-by-strain mutation count matrix for each transfer regime and randomizing combinations of mutation counts, constrained on the total number of mutations acquired within each gene across strains and the number of mutations acquired within each strain. Randomization was performed for 10,000 iterations using a Python implementation of the ASA159 algorithm. A null distribution was obtained using this approach, from which observed values of 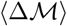 were standardized 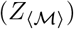 [46, 47]. To determine whether spore-forming genes and the *spo0A* regulon were enriched we calculated the difference in the fraction of nonsynonymous mutations between the WT and Δ*spo0A* strains for a given transfer regime, where the null was obtained using the binomial distribution in Eq. 9. Genes in the *spo0A* regulon were identified using a published database [48] and genes associated with endospore formation were identified by annotation. Finally, we identified genes that were preferentially enriched in one background or transfer regime over the other using the Skellam distribution as previously described [49].

### Simulating evolution with a seed bank

To determine whether the empirical patterns of genetic diversity we observed were consistent with the outcomes predicted by seed bank effects, we performed forward-time simulations based on a modified standard master equation that captures the joint evolution between active and dormant individuals [50]. Dormancy was incorporated in a way similar to previous research efforts [19, 21, 51, 52]. We focused on modeling the probability that out of a total of *A* active individuals that there are *a* individuals with fitness *X* at time *t* + *dt* (*p_A_*(*a*, *X*, *t* + *dt*)). Likewise, we are interested in the analogous probability for the pool of *D* dormant individuals, *p_D_*(*d*, *X*, *t* + *dt*). The selection coefficient of an individual (*s*) is a measure that is relative to the mean fitness of the population at the current generation 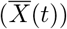, corresponding to a birth rate of 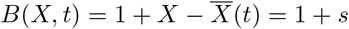 and a death rate of 1 for active individuals. We then calculated the probabilities of birth and death events among active individuals with fitness *X* that did not enter the seed bank on an interval of time of *dt* as

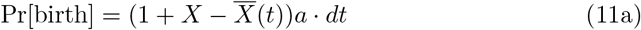

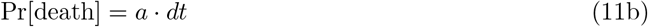

We then turn to the effect of mutation. Active individuals acquire mutations via reproduction, the fitness effects of which can be viewed as samples from a distribution of fitness effects (*ρ*(*s*)). Given that the rapid increase in allele frequencies we observed were likely to be predominantly driven by positive selection, we focused on a distribution of fitness effects of beneficial mutations with beneficial mutations occurring at a per-individual rate of *U_b_*. Taking the integral to account for *ρ*(*s*) as a continuous distribution, we obtained the following probability

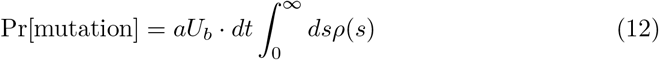

Finally, it is necessary to incorporate transition probabilities for dormancy. Given that a population consists of *A* active and *D* dormant individuals, we modeled dormancy by considering the number of individuals that enter or exit a dormant state each generation, *c*, chosen so that population sizes remain constant over time. Given *c*, an active individual enters a dormant state with probability 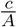 and a dormant individual resuscitates with probability 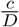. From this, we obtained the following transition probabilities

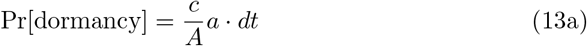

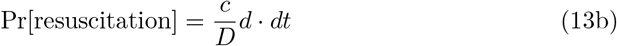

The length of time that an individual remains in a dormant state is a geometrically distributed random variable, where the probability of resuscitation (i.e., success probability) is the inverse of the per-individual probability of resuscitation. Using this distribution, the impact of dormancy is captured by the mean number of generations that an individual spends in a dormant state, 〈*T_d_*〉 = *D/c* and the ratio of active and dormant individuals *K* = *A/D*.

Using these transition probabilities, we defined the complement probability that none of them happen as Pr[nothing] such that the total probability sums to one. After defining equivalent transition probabilities for the dormant pool, we obtained

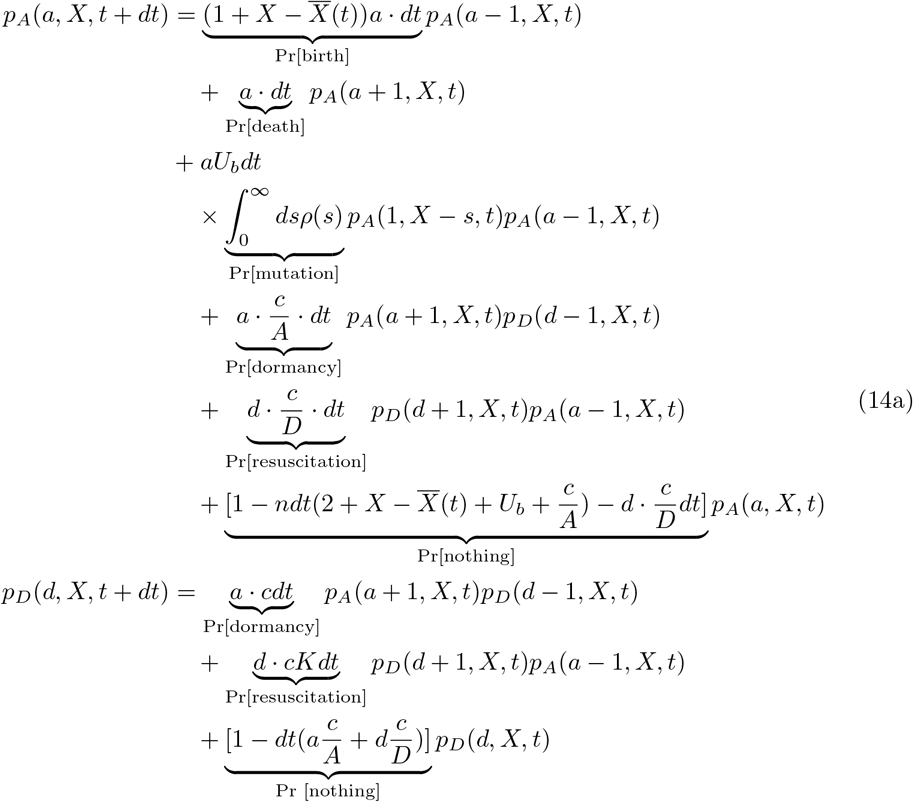

where Pr[nothing] is the probability that nothing happens and is calculated by adding all the transition probabilities and subtracting it from unity. The death rate of dormant individuals is likely much smaller than that of active individuals, if not nonexistent, allowing us to omit them from *p_d_*(*d*, *X*, *t* + *dt*). To maintain a constant population size, there needs to be an individual with a fitness value of *X* – *s* in order to acquire a *de novo* mutation with fitness s to increase the number of individuals with fitness *X* by one.

While the true value of *U_b_* is unknown, we elected for a value that was on the order of magnitude of 10% of the total mutation rate (all non-lethal mutations) obtained from a previously published mutation accumulation experiment *U_b_* = 10^-4^/indiv. [53]. The form and parameters of *ρ*(*s*) are virtually never known and difficult to infer on evolution experiments with timescales < 10^4^ generations [54]. In light of this fact, we elected to model *ρ*(*s*) as an exponential distribution with a scale parameter of 10^-2^. Simulations were run for 3,300 generations with *A* = 10^6^ for *c* = 10^-5^ and values of *D* ranging from 10^1^ — 10^6^. Only values of D were manipulated as the same transition rates can be obtained by manipulating *c*. Ten replicate simulations were performed for each value of D. All simulations were performed using custom Python scripts.

## Results and Discussion

By reconstructing the molecular fossil record of our experiment, we examined the trajectories of *de novo* mutations for all replicate populations (Fig. 1). The molecular evolutionary dynamics of *B. subtilis* largely followed our predictions, as the presence of a seed bank increased the maximum amount of genetic diversity retained by a population and decreased its per-generation rate of accumulation (Fig. 2; Table 1). Measures that capture distinct features of mutation trajectories were also largely consistent with our predictions (Fig. 3), results that were validated with forward-time population genetic simulations (Fig. 4). By comparing gene mutation counts between strain-transfer combinations (Fig. 5,6; Table 2), we determined that the ability to enter a dormant state radically altered the targets of molecular evolution, though the strength and direction of the effect was environment-dependent. The results of this long-term experiment test, confirm, and challenge long-standing hypotheses regarding the effect of seed banks on the dynamics of molecular evolution.

**Figure 1.**
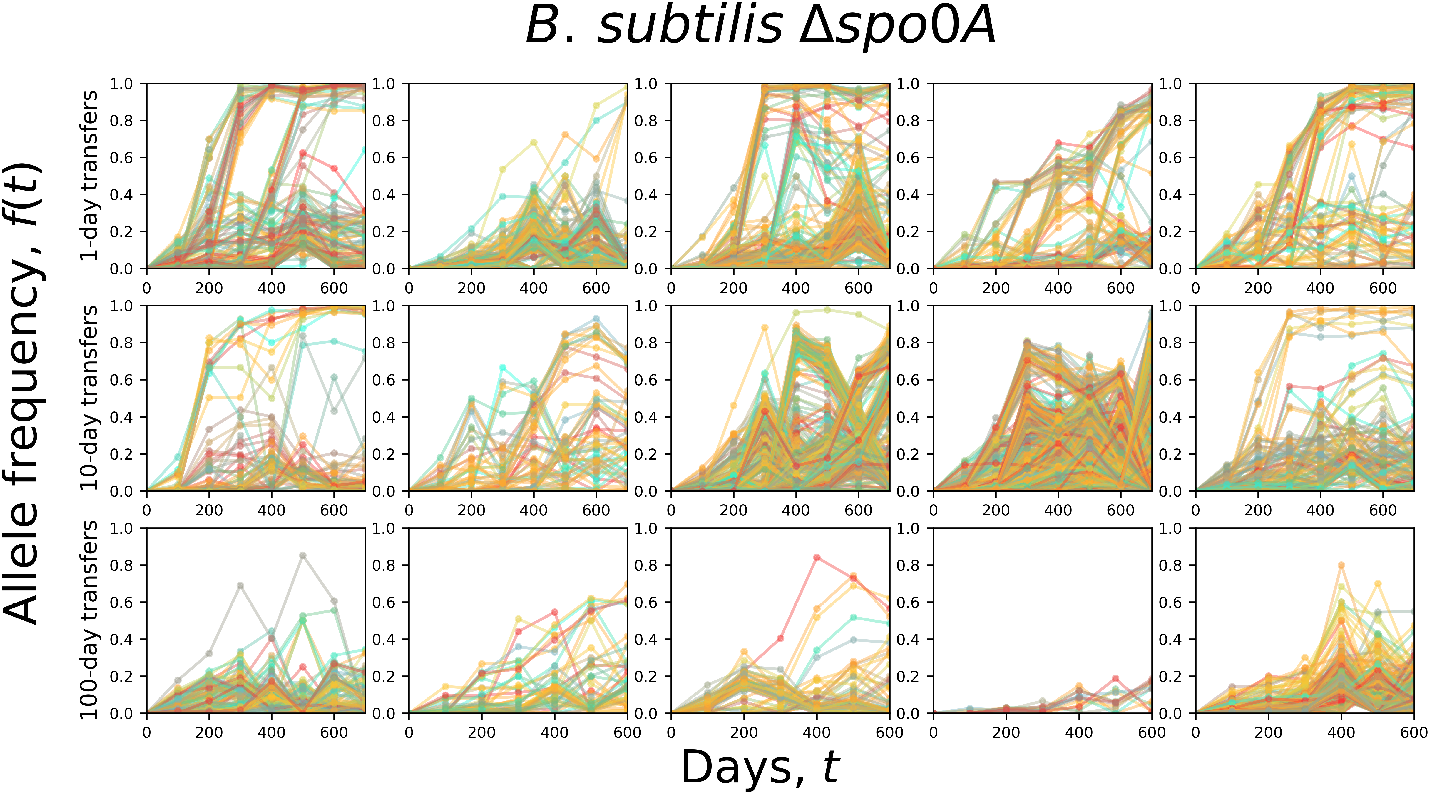
The allele frequency trajectories of *Bacillus subtilis* populations that were unable to form seed banks (Δ*spo0A*) exhibited qualitatively different dynamics across transfer regimes. Typically, a lower number of mutations and fixation events accumulated in the 10-day regime relative to the 1-day regime. There were even fewer detectable mutations in the 100-day regime, a likely consequence of the low number of generations that occurred over 700 days. All Δ*spo0A* replicate populations are included on this plot.

**Figure 2.**
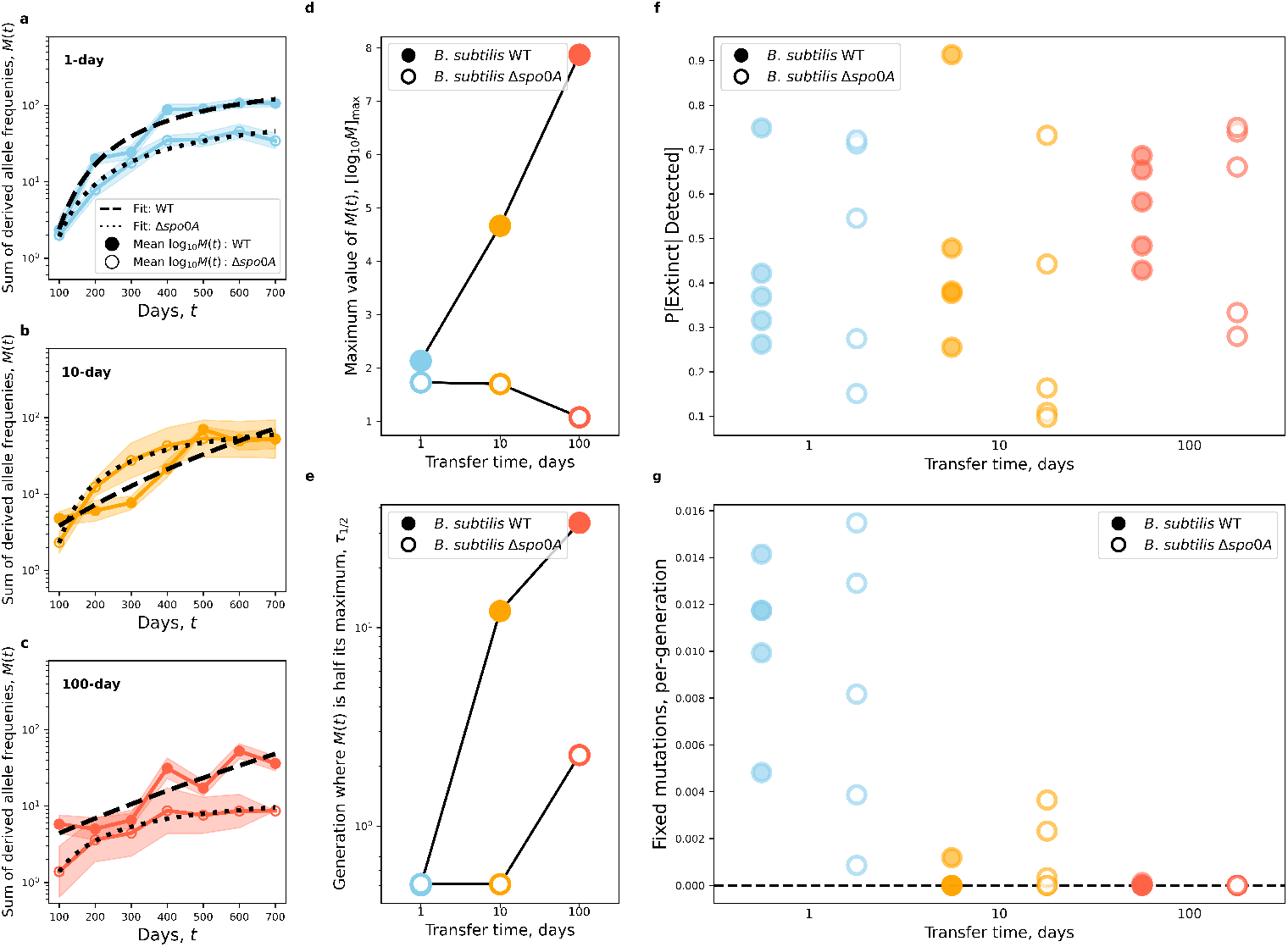
The presence of a seed bank altered the accumulation of genetic diversity. **a-c)** By examining the sum of derived allele frequencies (*M*(*t*), Eq. 2), we were able to summarize the accumulation of *de novo* mutations over time for all strains and transfer regimes. The WT strain had higher a mean log_10_*M*(*t*) than Δ*spo0A* and the relationship between *t* and log_10_*M*(*t*) became noticeably more linear as transfer time increased for the WT strain. Shaded areas represent the standard error of mean log_10_*M*(*t*). **d,e)** To quantify the effect of seed bank formation on this empirical relationship we formulated a phenomenological model that allowed us to summarize the curve through two parameters: the maximum amount of genetic diversity that could accumulate ([log_10_*M*]_max_) and the number of generations until half of the maximum is reached (*τ*_1/2_; Eq. 3). Values of [log_10_*M*]_max_ for the WT strain steadily increased with transfer time while Δ*spo0A* remained consistent with the prediction that the presence of a seed bank increases the amount of genetic diversity that a population can maintain. Conversely, *τ*_1/2_ decreased for Δ*spo0A* but remained constant for the WT, consistent with the prediction that the rate of molecular evolution would increase in the absence of a seed bank. **f,g)** However, the effect of seed banks on the final states of alleles was less straightforward. While fixation events occurred across transfer regimes and strains, there was substantial variation across replicate populations that made it difficult to determine whether the presence of a seed bank affected the probability of fixation or the rate of molecular evolution (per-generation number of substitutions).

**Figure 3.**
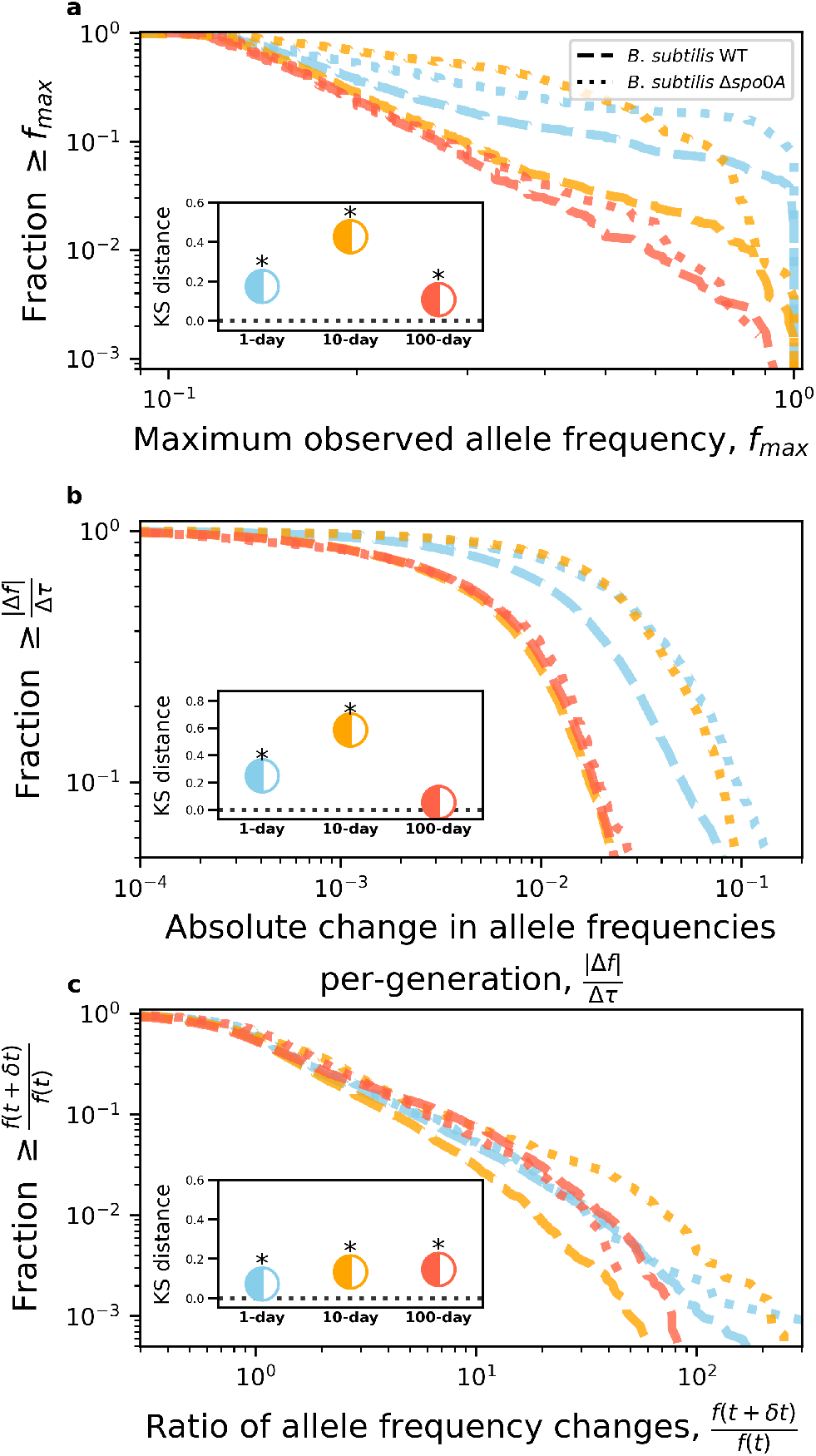
Due to the low number of fixation events we devised alternative measures of molecular evolution to evaluate the effect of seed banks. We examined three measures corresponding to **a)** the maximum frequency realized by an allele, **b)** the per-generation magnitude of change in allele frequency, and **c)** the change in the direction of allele frequencies between time points (Eq. 14). These measures were examined by calculating the empirical survival distribution (the complement of the empirical cumulative distribution function) for a given strain-transfer combination. The typical value of all three measures was higher for Δ*spo0a* than the WT across transfer regimes, consistent with the predicted effect of a seed bank. The difference we observed between strains was confirmed via Kolmogorov–Smirnov tests for all transfer regimes (Benjamini-Hochberg corrected *P*-values < 0.05 marked by asterisk).

**Figure 4.**
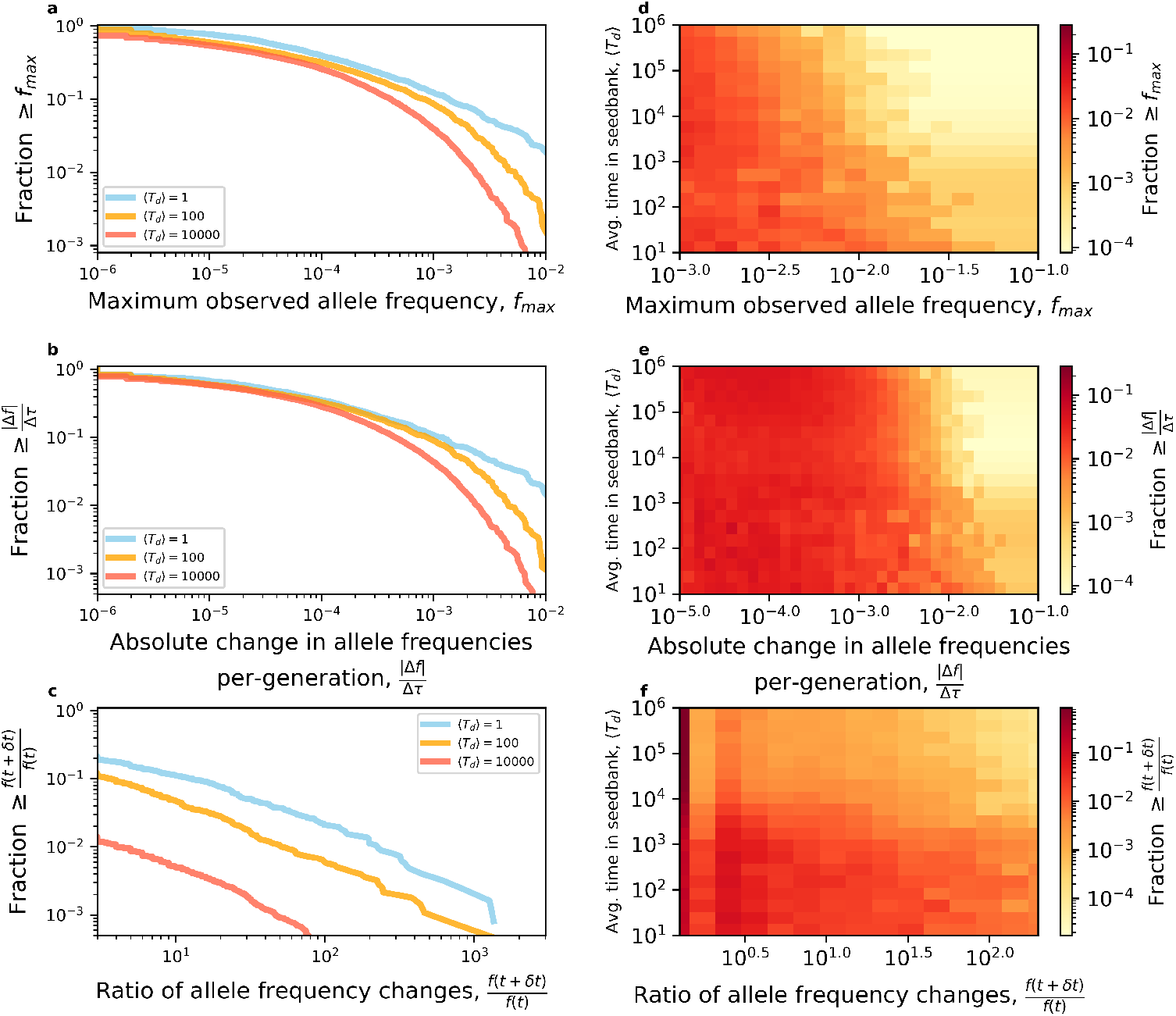
To validate our empirical observations we performed forward-time population genetic simulations of populations with seed banks of varying size, where the effect of seed banks can be summarized by the average number of generations that an individual spends in a dormant state (〈*T_d_*〉; Materials and Methods). **a)** The breadth of simulated survival distributions gradually decreased with 〈*T_d_*〉 for all three measures of allele frequencies (Eq. 14), recapitulating the empirical results described in Fig. 3. **b)** The robustness of this pattern is made evident by performing simulations across a wide range of 〈*T_d_*〉 values.

**Figure 5.**
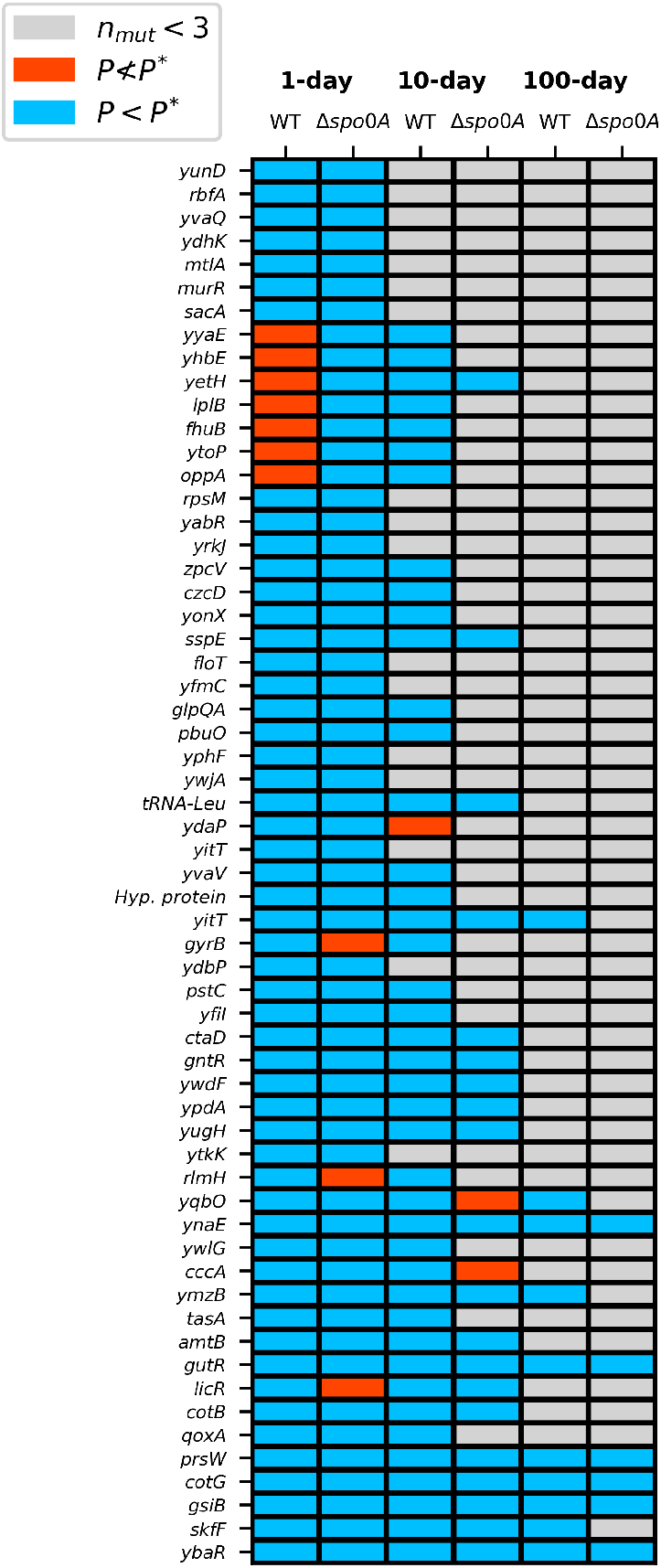
By comparing the set of genes that contributed towards parallel evolution with a given strain-transfer combination, we visualize patterns of convergent and divergent molecular evolution. While all strain-transfer combinations consistently acquired more nonsynonymous mutations than expected by chance at a large number of genes (*P* < *P*\*), very few genes were enriched exclusively within a given strain. Those genes that were significantly enriched within a given strain combination typically also acquired mutations at a non-significant level (*P* ≮ *P*\*) in the remaining strain, suggesting that the removal of endospore formation did not generate evolutionary trajectories that were divergent in terms of gene identity. To increase statistical power we ignored all genes that acquired less than three mutations (*n_mut_* < 3) across all five replicate populations for a given strain-transfer regime combination. Gene names are listed as provided in the annotated reference genome; all other names were acquired using RefSeq IDs (see File S1 for gene metadata).

**Figure 6.**
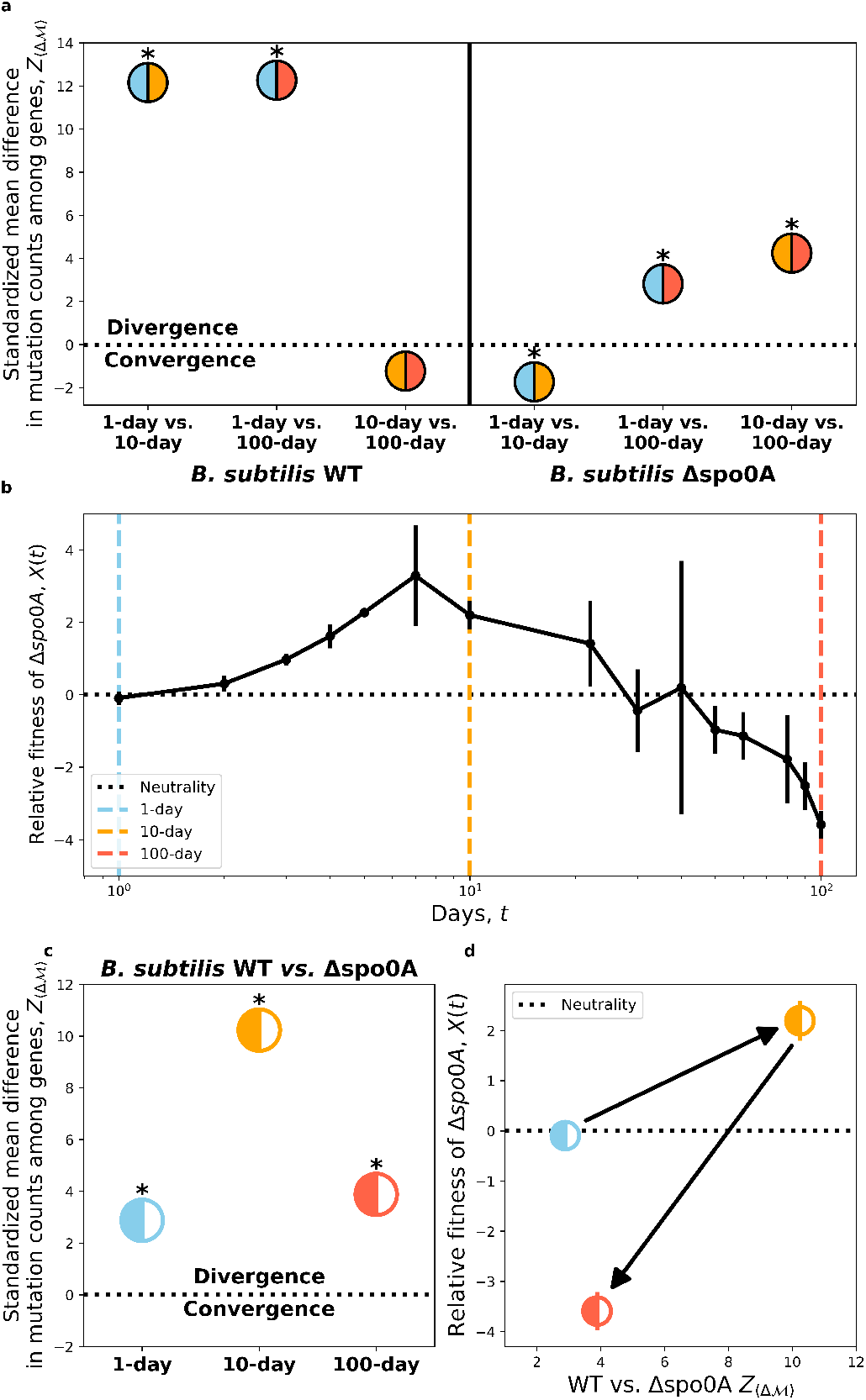
By examining the number of mutations within each gene, we determined whether convergent or divergent evolution occurred between a given pair of transfer regimes or strains by calculating the mean absolute difference of mutation counts across genes (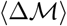; Eq. 10). **a)** A comparison between all transfer regime combinations within each strain reveals contrasting dynamics of divergence/convergence. Divergent evolution initially occurred among the WT background initially for 1-dat vs. 10-day and 1-day vs. 100-day comparisons, with 10-day and 100-day transfers having a weak signal of convergence. This pattern became inverted for Δ*spo0A*, as 1-day and 10-day transfers converged while the remaining transfer regime combinations diverged. For comparisons between strains, we compared signals of convergent/divergent evolution between WT and Δ*spo0A* strains with the transfer regime-dependent fitness effects of removing *spo0A*. **b)** A 100-day fitness assay between Δ*spo0A* and WT strains revealed the time-dependent fitness effect of the ability to form endospores. In the 1-day regime there is no fitness effect of *spo0A* removal, which ultimately becomes beneficial by day 10. However, shortly thereafter it became highly deleterious. Black dots represent the mean of three replicate assays while bars represent the standard error. **c)** Divergent evolution consistently occurred across transfer regimes, with the 10-day transfers harboring the strongest signal of divergent evolution. **d)** Mapping signals of divergent evolution to estimates fitness, we determined how the sign and magnitude of selection changed with the degree of divergent evolution. Asterisks denote *P* < 0.05.

**Table 1.**
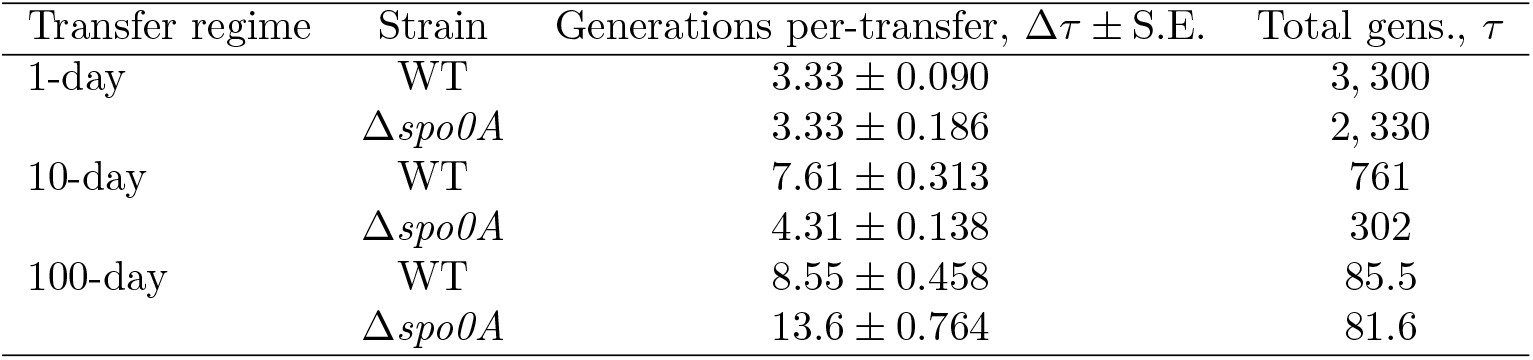
To evaluate the evolutionary effects of seed bank formation we maintained replicate populations of endospore forming (WT) and non-forming (Δ*spo0A*) strains of *B. subtilis* in environments with increasing degrees of energy-limitation, conditions where the ability to enter a dormant state is adaptive. To manipulate energy-limitation, we extended the time between resource replenishment via serial transfer, resulting in transfer regimes of 1, 10, and 100 days. The population size declined at different rates over time for WT and Δ*spo0A* strains in the 10 and 100-day transfer regimes, implying that the number of cells passaged between transfers varied across strain-transfer regime combinations. Under a standard serial dilution regime, the number of generations was estimated as the binary logarithm of the final population size after a transfer (*N_f_*) divided by its initial size (*N_i_*). By obtaining estimates of *N_i_* for each strain-transfer regime combination, we obtained a more accurate estimate of the number of generations that accrued per-transfer (Δ*τ*). Using these per-generation estimates we calculated the total number of generations (*τ*), a quantity that represents the evolutionary timescale of the experiment.

| Transfer regime | Strain | Generations per-transfer, $\Delta\tau \pm \text{S.E.}$ | Total gens., $\tau$ |
| --- | --- | --- | --- |
| 1-day | WT | $3.33 \pm 0.090$ | 3,300 |
| | $\Delta spo0A$ | $3.33 \pm 0.186$ | 2,330 |
| 10-day | WT | $7.61 \pm 0.313$ | 761 |
| | $\Delta spo0A$ | $4.31 \pm 0.138$ | 302 |
| 100-day | WT | $8.55 \pm 0.458$ | 85.5 |
| | $\Delta spo0A$ | $13.6 \pm 0.764$ | 81.6 |

**Table 2.** The number of genes that were enriched for nonsynonymous mutation solely within a given strain-transfer regime combination (i.e., unique genes).

| Transfer regime | Number unique enriched genes |  |
| --- | --- | --- |
| | *B. subtilis* WT | *B. subtilis* $\Delta spo0A$ |
| 1-day | 77 | 2 |
| 10-day | 12 | 8 |
| 100-day | 12 | 0 |

### The demographic effect of seed banks in rapidly evolving populations

While Δ*spo0A* populations were unable to form endospores, the fraction of endospores varied across transfer regimes for WT lines, providing the mean to assess the impact of seed bank formation on molecular evolutionary dynamics. Endospores were not detected in the 1-day transfer regime for the WT, though by the end of 10 days the proportion of spores reached 0.65 ± 0.062 (mean ± standard error). This estimate of the fraction of spores was consistent with prior energy-limitation experiments with *B. subtilis* [28].

Despite the inability to form protective structures, all replicate populations of Δ*spo0A* from all transfer regimes survived the experiment. This pattern of consistent demographic survival is a marked difference from a previous experiment that used an identical energy-limitation design, where bacterial taxa from diverse phyla (Bacteroidetes, Proteobacteria, and Deinococcus-Thermus) frequently underwent demographic extinction (i.e., *N* = 0) in 10 and 100-day transfer regimes [35], whereas *Bacillus* WT never went extinct. This comparative result suggests that *Bacillus* is exceptionally capable of surviving and evolving in harsh environments, even without access to what is generally considered its primary survival strategy of endospore formation.

### Seed banks altered the accumulation, but not the fates, of *de novo* mutations

Through pooled population sequencing, we reconstructed the trajectories of *de novo* mutations for all five replicate populations from each transfer regimes (Fig. 1; S2). By examining the sum of derived allele frequencies at a point in time (*M*(*t*)) we were able to examine how *de novo* mutations accumulated over the course of the experiment for all replicate populations (Fig. 2a-c, S3). Populations capable of forming seed banks tended to have higher value of *M*(*t*) in the 1 and 100-day transfer regimes, though this difference is not as apparent for the 10-day regime.

These patterns spurred the development of a novel mathematical model. Intuitively, seed banks can affect the accumulation of *de novo* genetic diversity over time through two means: 1) increasing the maximum amount of diversity that a populating can retain and 2) reducing the rate that diversity accumulates. Building off of this assessment, we developed a phenomenological model with two intuitive parameters (Eq. 3) that can be used to test our predictions, the maximum value of the logarithm of *M*(*t*) ([log_10_*M*]_max_) and the number of generations required for log_10_*M* to reach half of its maximum value (*τ*_1/2_). Estimates of these parameters obtained through numerical optimization and estimates of the number of generations that accrued within each treatment (Table 1) revealed that [log_10_*M*]_max_ remained fairly constant as energy-limitation (i.e., time between transfers) increased for the WT, but sharply decreased for Δ*spo0A* (Fig. 2d). In contrast, *τ*_1/2_ remained constant for Δ*spo0A*, but decreased as transfer time increased (Fig. 2e). This pattern was consistent with our prediction that the rate of accumulation of genetic diversity will be higher in populations that cannot form a seed bank.

While the trends we observed were generally robust, the error in our estimates for the 100-day transfer regime was considerable (Table S2). This error was likely a result of the small number of mutations acquired among populations in the 100-day treatment as well as their small population sizes, increasing the variance in our estimates of *M*(*t*). Regardless, parameter estimates consistently change in directions predicted by the seed bank effect.

Given that seed banks altered the accumulation of genetic diversity, we then examined how the presence of a seed bank altered the fate of a given mutation. The presence of seed banks reduced the efficiency of selection and retain genetic diversity [17], suggesting that a given mutation would have a lower probability of extinction as well as a lower substitution rate [18, 19]. Our estimates of the probability of extinction for a given mutation provide little evidence to support this claim, as there was substantial variation across replicate populations for a given strain-transfer combination (Fig. 2f). This result suggests that seed banks primarily altered the accumulation of genetic diversity by buffering the dynamics of segregating mutations rather than reducing their rate of extinction. However, we note that this study, as is typical of microbial pooled sequencing studies, has a lower frequency resolution 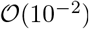. Given the large empirical sizes of our experimental populations, we were unable to evaluate the effect of dormancy on the evolutionary fates of mutations across a wide range of frequency values (1/*N* < *f* ≲ 10^-2^).

While the rate of fixation was similar for WT and Δ*spo0A* in the 1-day regime, as expected given that the ability to form a seed bank does not contribute to survival on that time scale, few fixation events occurred within replicate populations in the 10-day and 100-day transfer regimes (Fig. 2g). This paucity of fixations meant that the substitution rate could not be estimated for certain transfer regimes, preventing comparisons from being performed. While the strength and direction of selection could be evaluated by examining the proportion of nonsynonymous to synonymous mutations (*pN/pS*), estimates of *pN/pS* were consistently less than one across transfer regimes and there was no difference in *pN/pS* values between WT and Δ*spo0A* populations within a given transfer regime (Fig. S4). This lack of difference suggests that purifying selection was predominant regardless of whether a population could enter the seed bank or in environments where the ability to enter a dormant state would be favorable. This consistent lack of difference is unlikely to be the result of our choice of knockout gene, as genome-wide signals of *pN/pS* were not affected by the exclusion of genes in the *spo0A* regulon (Table S3).

### Seed banks altered the dynamics of segregating mutations

While seed banks had a clear effect on the accumulation of *de novo* genetic diversity, they did not substantially alter the evolutionary fates of mutations. In addition, few if any fixation events were inferred for 10 and 100-day transfer regimes. These two results indicate that the substitution rate and the probability of extinction of a mutation are uninformative for this particular study. Such quantities are useful in that their value in the presence of a seed bank can be compared to predictions from existing mathematical models [18, 19, 21]. However, their emphasis on the final state of a given mutation (i.e., fixation or extinction) means that these quantities do not necessarily capture the dynamics mutations exhibit over their sojourn times. Therefore, we leveraged the temporal structure of the data by examining mutation trajectories in order to evaluate the extent that seed banks alter the dynamics of molecular evolution.

We examined three measures of each mutation trajectory: 1) the maximum frequency that a mutation reached (*f_max_*), 2) the set of per-generation magnitudes of frequency changes 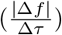 over time, and 3) the set of changes in the direction of allele frequency changes between observations over time (*f* (*τ* + Δ*τ*)/*f* (*τ*)). By calculating empirical survival distributions for a given measure for each strain-treatment combination, we determined the degree that seed banks altered the molecular evolutionary dynamics of *B. subtilis*.

First, given the dearth of fixation events, it is worth investigating whether the maximum frequency that a mutation obtained over its sojourn time (*f_max_*) is a sufficient proxy of fixation. The condition that a mutation reached a sufficiently high frequency guarantees fixation under a single-locus model of evolution [55]. Though this certainty is not necessarily the case when multiple beneficial mutations simultaneously segregate, where recombination may not be sufficiently high to justify the assumption of quasi-linkage equilibrium (i.e., independent evolution among multiple sites). Regardless, there are various derivations in theoretical population genetics indicating that *f_max_* is reflective of the strength and direction of selection as well as the conditional probability of fixation of an allele [56–58].

We do not infer fixation events in several replicate populations, so we cannot confirm that there is a positive empirical relationship between *f_max_* and the probability of fixation. However, we inferred a large number of mutation extinction events across all replicate populations. Given that a segregating allele has to eventually become extinct of fixed, the probability of fixation can be viewed as the complement of the probability of extinction (Pr[fixation =1 – Pr[extinction]), meaning that if *f_max_* is reflective of fixation then we should observe an inverse relationship between estimates of Pr[extinction|detected] and *f_max_*. We find that this prediction holds, as all strain-transfer regime combinations exhibit the inverse relationship (Fig. S5), allowing us to proceed with our analyses of the distribution of *f_max_*.

Across transfer regimes, Δ*spo0A* populations had higher values of *f_max_* than their corresponding WT transfer regime and the significance of the distance between the distributions was tested and confirmed via Kolmogorov–Smirnov tests (Fig. 3a). Interestingly, WT and Δ*spo0A* populations had the smallest distance in the 100-day transfer regimes, which is likely due to the comparatively brief evolutionary timescale reducing the maximum attainable frequency of a *de novo* mutation. Overall, the values of *f_max_* exhibited in the presence and absence of a seed bank are consistent with the prediction that the presence of a seed bank will reduce the rate of molecular evolution.

The presence of a seed bank generated a similar effect on the magnitude of allele frequency changes between observations 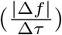 and the direction of frequency changes (*f*(*τ* + Δ*τ*)/*f*(*τ*); Fig. 3b,c). There was a small, but significant difference where Δ*spo0A* had a higher magnitude of change. This distance increased considerably for the 10-day transfer regime, consistent with the prediction that the presence of a seed bank would buffer temporal changes in allele frequencies. Though much of this distance disappeared for the 100-day transfer regime, again, a likely result of the comparatively brief evolutionary timescale of the treatment.

While the distributions of quantities calculated from mutation trajectories matches our predictions regarding the effect of seed banks on molecular evolutionary dynamics, they are population genetic quantities that are not typically examined. Arguably this reflects the historic difficulty of obtaining temporally resolved frequency trajectories for a large number of mutations, rather than the measures themselves being uninformative. To evaluate how these measures of mutation trajectories typically behave in the presence of a seed bank, we simulated a master equation describing an adapting population that incorporates seed bank dynamics using previously proposed mathematical formulations [19, 21] (Eq. 14; Fig. S6). Briefly, in a population with *A* active individuals *c* individuals enter a seed bank of size *D*, while a corresponding number of individuals are resuscitated from their dormant state. The effect of this dynamic can be captured by the average number of generations that an individual spends in a dormant state (〈*T_d_*〉 = *D/c*) and the ratio of active and dormant individuals *K* = *A/D* [19]. While c is difficult to estimate, *A* and *D* can readily be measured, providing empirical constraint on the range of parameter values in our simulations. Furthermore, endospores are undetectable in WT lines and absent entirely in Δ*spo0A* (i.e., *D* = 0), meaning that the difference between WT and Δ*spo0A* mutation trajectory statistics in Fig. 3 for 1 and 10-day regimes can be viewed as the degree that the ability to enter a dormant state decreased the typical value of a given mutation trajectory statistic. We captured this seed bank effect by performing simulations across a range of values of *c* for empirically informed values of *A* and *D* and estimating the measures in Eq. 4 from simulated trajectories. Using this approach, we were able to recapitulate our empirical survival distributions (Fig. 4). These principled simulations combined with empirical observations confirm long-standing untested hypotheses regarding the effect of seed banks dynamics of molecular evolution.

The Δ*spo0A* populations were incapable of forming endospores and exhibited mutation trajectories that were consistent with an absence of a seed bank. However, *spo0A* can regulate the genetic competence of *B. subtilis* [59–61], which in turn affects the rate of recombination. To determine whether Δ*spo0A* populations had a lower recombination rate, we calculated squared correlation coefficient between all pairs of segregating mutations within each replicate population. This measure has previously been used to examine the effect of recombination on rapidly evolving microbial populations [44]. By pooling coefficients across replicate populations and comparing the distributions of WT and Δ*spo0A* strains, we found that the degree of correlation was consistently greater for Δ*spo0A* across transfer regimes (Fig. S7a). This result suggests that the rate of recombination was lower in Δ*spo0A* populations relative to the WT. To determine whether this pattern was driven by the absence of a seed bank, we calculated distributions of correlation coefficients from simulated data obtained using Eq. 14 and found that seed banks alone were insufficient to cause the patterns we observed (Fig. S7b).

While the distribution of correlation coefficients suggests that the rate of recombination was lower in Δ*spo0A* populations, this does not prove that the statistical patterns we observed in Fig. 3 were caused by a difference in recombination rates between Δ*spo0A* and the WT. To determine whether recombination could feasibly be responsible for the distribution of single-locus measures we examined, we developed and implemented a novel simulation based on recent theoretical developments where two alleles on different loci segregate in a population with dormant and active individuals (Supplemental Information; Eq. S1) [58]. We found no evidence that the KS distance between dormancy capable and incapable populations was dependent on the population scaled recombination rate for all trajectory summary statistics (Fig. S8), suggesting that any purported difference in the recombination rate between WT and Δ*spo0A* would not be capable of generating the between-strain KS distance values estimated in the 10-day transfer regime. Thus, the analyses presented in Fig. 3 are reflective of the effect of seed banks formation.

### Parallelism and (con/di)vergence at the gene level

To determine whether endospore formation as a life-history strategy affects the targets of molecular evolution in addition to its direction, it was necessary to first identify the potential contributors of adaptation among replicate populations within a given strain-transfer combination. To identify potential contributors towards adaptation, we examined the distribution of nonsynonymous mutation counts across genes within each strain-transfer combination. The log-likelihood that some number of genes were enriched for nonsynonymous mutations (Δ*ℓ*) was significant across 1, 10, and 100-day transfer regimes for both strains (Table S4). However, values of Δ*ℓ* tended to be slightly elevated for the WT strain, suggesting that a higher degree of parallel evolution occurred when endospore formation was possible. To determine whether this difference between strains was real or an artifact of the higher number of mutations acquired by the WT strain across transfer regimes, we randomly sub-sampled mutations to obtain a distribution of Δ*ℓ* values for each strain-transfer combination. While the difference in Δ*ℓ* values between strains was greatly reduced, the WT strain consistently had a higher degree of genome-wide parallelism across transfer regimes (Fig. S9a). This increased degree of parallelism suggests that the presence of a seed bank can make the molecular targets of evolution more predictable. However, it is worth considering whether this result is due to the use of Δ*spo0A*. The gene *spo0A* is a master regulator that controls cellular processes in addition to endospore formation [62, 63], meaning that the decrease in parallelism in Δ*spo0A* could be a consequence of pleiotropy caused by the *spo0A* regulon. That the *spo0A* regulon is enriched for nonsynonymous mutations in the 1-day transfer regime (Table S5) supports this hypothesis. To investigate this possibility, we performed the same sub-sampling procedure on the set of genes that are not in the *spo0A* regulon and compared the sub-sampled likelihood to the result obtained by using all genes in the genome. We found the contribution of *spo0A* to the genome-wide signal of parallelism to be inconsequential, suggesting that the deletion of *spo0A* was not responsible for the patterns of parallel evolution we observed (Fig. S9b).

To deconstruct this genome-wide pattern of parallelism, we examined the excess number of nonsynonymous mutations at a given gene (i.e., multiplicity [40]). As expected, the distribution of multiplicities of the WT was consistently higher than Δ*spo0A* across transfer regimes (Fig. S10a-c), suggesting that this effect is independent of energy limitation and is instead likely driven by the larger number of nonsynonymous mutations that accrued in the WT strain. To account for this difference, we calculated a *P*-value for each gene within each strain-transfer combination, allowing us to pare down the total number of genes in the genome to a small number of genes that were disproportionately enriched for nonsynonymous mutations, the putative targets of selection. Identifying the set of significantly enriched genes revealed that genes enriched within the WT for a given transfer regime also tended to be enriched within Δ*spo0A* (Fig. 5, S10d-f). This pattern of consistent enrichment occurred across transfer regimes as well as among transfer regime comparisons within a given strain (Fig. S11,S12), suggesting that, generally, the direction of evolution at the gene-level tended towards convergence rather than divergence. We found that this is the case, as the degree of overlap in enriched genes relative to a null distribution suggests convergent evolution (Fig. S13 [35]). While certain transfer regime and strain comparisons had stronger signals of convergence than others, overall convergent evolution overwhelmingly occurred.

Similar to a previous analysis [35], it is likely that gene identity was, again, too coarse a measure to determine whether convergent or divergent evolution occurred. While Δ*spo0A* is a master regulatory gene, its removal may have only slightly perturbed the rates of evolution across a large number of genes in a given environment. If true, then it is arguably more appropriate to examine the difference in mutation counts among enriched genes in order to assess whether convergent or divergent evolution occurred. By examining the mean absolute difference in mutation counts across enriched genes between two transfer regimes 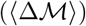 and standardizing the observed value with respect to a null distribution 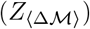 obtained via permutation, we established whether convergent or divergent evolution occurred. The WT strain exhibited significant divergent evolution for the 1-day vs. 10-day and 1-day vs. 100-day comparisons, a result that was consistent with the WT surviving energy-poor environments by forming endospores as a life-history strategy (Fig. 6a). This conclusion is strengthened by the evidence of convergent evolution for the 10-day vs. 100-day comparison, though it was ultimately not significant. For Δ*spo0A* the pattern changed in a manner that resembled a reflection of the WT pattern. There was a significant signal of convergent evolution for the 1-day vs. 10-day comparison though for the 1-day vs. 100-day and 10-day vs. 100-day comparisons we found a strong signal of divergent evolution. To summarize, the removal of Δ*spo0A* generated opposite trends in the direction of molecular evolution at the gene level across combinations of transfer regimes.

To examine how seed bank formation affected the direction of molecular evolution at the gene level within a given transfer regime, we repeated the convergent/divergent analysis between the WT and Δ*spo0A* populations, analyses that can be bolstered through comparisons to fitness estimates (Fig. 6b). Divergent evolution overwhelmingly occurred for all three comparisons, though it was at its highest for the 10-day transfer regime (Fig. 6c). This increased divergence for the 10-day transfer regime, along with evidence of convergent evolution for the 1-day vs. 10-day comparison, may reflect adaptation of Δ*spo0A* to this specific regime. Though, again, it is worth investigating the contribution of genes within the *spo0A* regulon, as the fraction of nonsynonymous mutations in the *spo0A* regulon was significantly higher in the Δ*spo0A* lines relative to the WT for the 10 and 100-day transfer regime (Table S6). However, it is unlikely that the pleiotropic nature of *spo0A* was responsible for the differing patterns of convergence and divergence observed between WT and Δ*spo0A* strains, as we obtained virtually indistinguishable estimates of 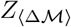 when genes in the *spo0A* regulon were excluded from our analysis (Fig. S14).

Our estimates of 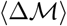 between strains for a given transfer-regime can be compared to the corresponding fitness effect of Δ*spo0A*, allowing us to examine how the sign and magnitude of fitness changes with the direction of molecular evolution. We found that after 1-day of co-culture that the fitness difference of Δ*spo0A* relative to the WT was undetectable via CFU counts, meaning that the fitness effect of being unable to form endospores was effectively zero. This result was consistent with the hypothesis that endospore formation has a negligible fitness effect in environments where it is unlikely to be expressed and, thus, would not be advantageous. However, we note that the removal of *spo0A* could have fitness effects in different environments that vary in other aspects but retain the regular resource replenishment of our experiment.

We found that the increase in fitness of Δ*spo0A* for the 10-day transfer regime was accompanied by an increase in the degree of divergent evolution (Fig. 6d). This result suggests that the ability to form endospores actually conferred a temporary fitness disadvantage at the 10-day mark. However, by day 100 the fitness benefit of Δ*spo0A* had dissipated and the magnitude of divergent evolution had diminished, suggesting that it is unlikely that Δ*spo0A* was able to adapt to the harsh 100-day environment. An analysis of the set of genes that were enriched for mutations within a specific strain-transfer combination supports this conclusion [49]. The 100-day Δ*spo0A* was the only strain-transfer combination with no unique enriched genes (Table 2; File S2), suggesting that Δ*spo0A* may have been unable to adapt to this extremely energy-limited environment.

While it is unlikely that Δ*spo0A* was able to adapt to the 100-day transfer regime, conversely, the 10-day Δ*spo0A* transfer regime harbored the highest number of unique enriched genes for all three Δ*spo0A* transfer regimes, suggesting that adaptation may have occurred even in the absence of endospore formation. The mechanism responsible for the temporary gain in fitness of Δ*spo0A* is unknown, though it is likely partially due to the recycling of dead cells, a phenotype that allows individuals to exploit an untapped resource [28, 38, 64]. Naturally, dormant cells cannot use this resource as their metabolism is effectively nonexistent and, in the case of endospores, metabolically inert, leaving Δ*spo0A* with unrestricted access. Regardless, for the purposes of this study, the removal of endospore formation as a life-history strategy provided a clear fitness benefit in certain energy-limited environments.

The disuse of a trait frequently results in a relaxation of selective pressure on its underlying loci [32, 33]. Given that endospore formation did not occur in the 1-day transfer regime, it is possible that the pathways encoding said life-history trait were susceptible to decay due to relaxed selective pressure. By calculating the fraction of nonsynonymous mutations in genes that encode for endospore formation and calculating the difference between Δ*spo0A* and the WT, we found that endospore encoding genes were slightly enriched in the WT for all transfer regimes, a difference that was significant using null distributions simulated via binomial sampling (Table S7). Operating under the premise that the majority of endospore-forming genes are nonfunctional in Δ*spo0A* populations, for WT populations in the daily transfer regime this result can be viewed as the outcome of positive selection for the removal of endospore formation as an energetically costly trait in energy-replete environments [65, 66]. Prior studies, as well as the spore accumulation assays we performed, support this conclusion [32–34]. The ability to form endospores became rapidly diminished for populations in the 1-day transfer regime over the first 500 days of the experiment, to to the point that it took 10 days for 10% of the population to form endospores (Fig. S15). However, the question remains as to why 100-day WT populations acquired a greater fraction of nonsynonymous mutations in endospore encoding genes. Endospore formation underwent no noticeable decline in the 100-day regime (Fig. S15), suggesting that these mutations had negligible or even positive effects.

## Conclusion

We demonstrated that the ability to form seed banks altered the molecular evolutionary dynamics of microbial populations. Populations capable of forming seed banks consistently accumulated higher levels of genetic diversity and had a reduced rate of molecular evolution. Through forward-time simulations, we were able to recapitulate empirical observations on the effect of seed banks on the rate and direction of allele frequency changes as well as the maximum attainable frequency. In addition to testing previously proposed predictions on the effect of seed banks on genetic diversity, new patterns were found. Specifically, we determined that endospore formation may have the capacity to alter the direction of molecular evolution within a population. Stated inversely, the absence of endospore formation contributed to a substantial signal of divergent evolution for populations in energy-limited environments. This signal of divergence, alongside the observation that the absence of endospore formation provided a substantial fitness benefit, suggests that adaptation to energy-limited environments may be possible in the absence of a highly conserved life-history strategy. Though any such adaptation would likely be transitory, as the absence of endospore formation resulted in an increasingly strong fitness disadvantage as the degree of energy-limitation increased.

While our study offers evidence that the ability to form a seed bank alters the rate and direction of molecular evolution in *B. subtilis*, *spo0A* remains a confounding variable. The gene is a master regulator that can have pleiotropic effects on biofilm formation [67–69], motility [70, 71], and genetic competence [59, 60, 72]. While we did not directly examine biofilm formation or motility, it is unlikely that they contributed towards adaptation as all replicate populations were maintained in well-aerated flasks in a constant shaker. However, the strength of correlations between mutation frequency trajectories remained higher in Δ*spo0A* across transfer regimes, a statistical pattern which suggests a lower rate of recombination (Fig. S7). Such a reduction could occur if *spo0A* played a major role regulating genetic competence in our environmental conditions. Though the rate of recombination did not alter the distance between simulated distributions of mutation trajectory statistics for dormancy capable and incapable populations (Fig. S8). Finally, the main conclusions of our analyses of parallelism and divergent/convergent evolution were not affected by the inclusion of the *spo0A* regulon (Fig. S9, S14), suggesting that the pleiotropic effects of the gene were not so considerable that they were able alter the distribution of mutation counts across genes or the degree of purifying selection.

This study tested lost-standing predictions and generated new questions for future research. Given that *Bacillus* incurs an environment-dependent fitness effect when endospore formation is removed, it is worth investigating the quantitative effects of mutation on endospore formation and, ultimately their effect on fitness. Given that limited evolutionary timescales can be obtained in experiments that mimic the environments where the ability to enter a dormant state has the greatest fitness benefit, alternative approaches may be necessary to probe the effect of mutation on endospore formation. One promising option is mass barcoding (e.g., RB-TnSeq), sidestepping the slow input of mutations via slow generation times by generating large mutant libraries, the frequencies of which can then be tracked [73]. Such approaches have already been applied to spores in members of the genus *Streptomyces* [74] and could be leveraged to identify environment-dependent fitness effects as well as the mode in which traits, such as endospore formation, mediate fitness effects in *Bacillus* [75]. Given that endospore formation is a complex trait with many loci that are likely interacting, it may be a suitable candidate to apply recently developed models that predict the form of the distribution of fitness effects when interactions between loci are prevalent [76].

## Data and code availability

Raw sequence data of Δ*spo0A* lines are available on the NCBI Sequencing Read Archive under BioProject ID PRJNA639642. Raw sequence data for WT lines was previously published [35] and is available under BioProject ID PRJNA639414. Reproducible code to perform all simulations and analyses is available on GitHub under the repository: Bacillus_Evol_Timeseries. Processed data and annotations are available on Zenodo under the DOI: 10.5281/zenodo.5549311.

## Acknowledgements

We thank P. G. Wall for providing a list of genes in the *spo0A* regulon. We thank M. Behringer, T. Doak, D. A. Drummond, P. Foster, B. K. Lehmkuhl, M. Lynch, J. McKinlay, and members of the Lennon lab for their helpful feedback at various stages of the project. We thank A. G. Casanova, D. A. Schwartz, and P. G. Wall for feedback on initial drafts. We thank H. Long for his help in troubleshooting library construction, K. Miller for assisting with sample collection, and members of the Kearns lab for sharing their plasmid protocols and assisting with troubleshooting. This work was supported by the US Army Research Office (W911NF-14-1-0411 to JTL), the National Science Foundation (DEB-1442246, DEB-1934554, DBI-2022049 to JTL), the National Aeronautics and Space Administration (80NSSC20K0618 to JTL) and the Society for the Study of Evolution Graduate Research Excellent Grant (GREG) Rosemary Grant Advanced Award (to WRS). Computing resources for simulations was supported by Lilly Endowment, Inc., through its support for the Indiana University Pervasive Technology Institute, the National Science Foundation under Grant No. CNS-0521433, and Shared University Research grants from IBM, Inc., to Indiana University.

## Author contributions

W.R.S., E.P., and J.T.L. designed the project; W.R.S., E.P., and K.B.G. conducted the experiments and generated the data; W.R.S. built and implemented the model; W.R.S. performed all analyses; W.R.S., E.P., K.B.G., and J.T.L. wrote the manuscript.

## References

1. Lennon, J. T. & Jones, S. E. Microbial seed banks: the ecological and evolutionary implications of dormancy. Nature Reviews. Microbiology 9, 119–130 (2011).

2. Buoro, M. & Carlson, S. M. Life-history syndromes: Integrating dispersal through space and time. Ecology Letters 17, 756–767 (2014). URL https://onlinelibrary.wiley.com/doi/abs/10.1111/ele.12275.

3. Rubio de Casas, R., Donohue, K., Venable, D. L. & Cheptou, P.-O. Gene-flow through space and time: dispersal, dormancy and adaptation to changing environments. Evolutionary Ecology 29, 813–831 (2015). URL https://doi.org/10.1007/s10682-015-9791-6.

4. Vitalis, R., Rousset, F., Kobayashi, Y., Olivieri, I. & Gandon, S. The joint evolution of dispersal and dormancy in a metapopulation with local extinctions and kin competition. Evolution 67, 1676–1691 (2013). URL https://onlinelibrary.wiley.com/doi/abs/10.1111/evo.12069.

5. Venable, D. L. & Brown, J. S. The selective interactions of dispersal, dormancy, and seed size as adaptations for reducing risk in variable environments. The American Naturalist 131, 360–384 (1988). URL https://www.journals.uchicago.edu/doi/abs/10.1086/284795.

6. Venable, D. L. & Lawlor, L. Delayed germination and dispersal in desert annuals: Escape in space and time. Oecologia 46, 272–282 (1980). URL https://doi.org/10.1007/BF00540137.

7. Jones, S. E. & Lennon, J. T. Dormancy contributes to the maintenance of microbial diversity. Proceedings of the National Academy of Sciences 107, 5881–5886 (2010). URL https://www.pnas.org/content/107/13/5881.

8. Tellier, A., Laurent, S. J. Y., Lainer, H., Pavlidis, P. & Stephan, W. Inference of seed bank parameters in two wild tomato species using ecological and genetic data. Proceedings of the National Academy of Sciences 108, 17052–17057 (2011). URL https://www.pnas.org/content/108/41/17052. Publisher: National Academy of Sciences Section: Biological Sciences.

9. Hairston, N. G. et al. Rapid evolution revealed by dormant eggs. Nature 401, 446–446 (1999). URL https://www.nature.com/articles/46731. Number: 6752 Publisher: Nature Publishing Group.

10. Huang, X. et al. The earliest stages of adaptation in an experimental plant population: strong selection on QTLS for seed dormancy. Molecular Ecology 19, 1335–1351 (2010). URL https://onlinelibrary.wiley.com/doi/abs/10.1111/j.1365-294X.2010.04557.x. _eprint: https://onlinelibrary.wiley.com/doi/pdf/10.1111/j.1365-294X.2010.04557.x.

11. Shoemaker, W. R. & Lennon, J. T. Evolution with a seed bank: The population genetic consequences of microbial dormancy. Evolutionary Applications 11, 60–75 (2018). URL https://www.ncbi.nlm.nih.gov/pmc/articles/PMC5748526/.

12. Guppy, M. & Withers, P. Metabolic depression in animals: physiological perspectives and biochemical generalizations. Biological Reviews of the Cambridge Philosophical Society 74, 1–40 (1999).

13. Willis, C. G. et al. The evolution of seed dormancy: environmental cues, evolutionary hubs, and diversification of the seed plants. New Phytologist 203, 300–309 (2014). URL https://nph.onlinelibrary.wiley.com/doi/abs/10.1111/nph.12782. _eprint: https://nph.onlinelibrary.wiley.com/doi/pdf/10.1111/nph.12782.

14. Doebeli, M., Ispolatov, Y. & Simon, B. Towards a mechanistic foundation of evolutionary theory. eLife 6, e23804 (2017). URL https://doi.org/10.7554/eLife.23804.

15. Kunkel, T. A. DNA replication fidelity. The Journal of Biological Chemistry 279, 16895–16898 (2004).

16. Lennon, J. T., den Hollander, F., Wilke-Berenguer, M. & Blath, J. Principles of seed banks and the emergence of complexity from dormancy. Nature Communications 12, 4807 (2021). URL https://www.nature.com/articles/s41467-021-24733-1. Bandiera_abtest: a Cc_license_type: cc_by Cg_type: Nature Research Journals Number: 1 Primary_atype: Reviews Publisher: Nature Publishing Group Subject_term: Applied mathematics;Biodiversity;Evolutionary theory;Theoretical ecology Subject_term_id: applied-mathematics;biodiversity;evolutionary-theory;theoretical-ecology.

17. Schoen, D. J., David, J. L. & Bataillon, T. M. Deleterious mutation accumulation and the regeneration of genetic resources. Proceedings of the National Academy of Sciences 95, 394–399 (1998). URL https://www.pnas.org/content/95/1/394. Publisher: National Academy of Sciences Section: Biological Sciences.

18. Koopmann, B., Müller, J., Tellier, A. & Živković, D. Fisher-Wright model with deterministic seed bank and selection. Theoretical Population Biology 114, 29–39 (2017).

19. Blath, J., Casanova, A. G., Eldon, B., Kurt, N. & Wilke-Berenguer, M. Genetic variability under the seedbank coalescent. Genetics 200, 921–934 (2015). URL https://www.genetics.org/content/200/3/921.

20. Kaj, I., Krone, S. M. & Lascoux, M. Coalescent theory for seed bank models. Journal of Applied Probability 38, 285–300 (2001). URL https://www.cambridge.org/core/journals/journal-of-applied-probability/article/coalescent-theory-for-seed-bank-models/3137A15F299546B0AA701FA974C19E46.

21. Blath, J., Casanova, A. G., Kurt, N. & Wilke-Berenguer, M. A new coalescent for seed-bank models. Annals of Applied Probability 26, 857–891 (2016). URL https://projecteuclid.org/euclid.aoap/1458651822.

22. van Gestel, J., Ackermann, M. & Wagner, A. Microbial life cycles link global modularity in regulation to mosaic evolution. Nature Ecology & Evolution 3, 1184–1196 (2019). URL http://www.nature.com/articles/s41559-019-0939-6.

23. Tellier, A. Persistent seed banking as eco-evolutionary determinant of plant nucleotide diversity: novel population genetics insights. New Phytologist 221, 725–730 (2019). URL https://onlinelibrary.wiley.com/doi/abs/10.1111/nph.15424. _eprint: https://onlinelibrary.wiley.com/doi/pdf/10.1111/nph.15424.

24. Heinrich, L., Müller, J., Tellier, A. & Živković, D. Effects of population- and seed bank size fluctuations on neutral evolution and efficacy of natural selection. Theoretical Population Biology 123, 45–69 (2018). URL https://www.sciencedirect.com/science/article/pii/S0040580917301715.

25. Nunney, L. The effective size of annual plant populations: The interaction of a seed bank with fluctuating population size in maintaining genetic variation. The American Naturalist 160, 195–204 (2002). URL https://www.journals.uchicago.edu/doi/full/10.1086/341017. Publisher: The University of Chicago Press.

26. Ellner, S. & Hairston, N. G. Role of overlapping generations in maintaining genetic variation in a fluctuating environment. The American Naturalist 143, 403–417 (1994). URL https://www.journals.uchicago.edu/doi/abs/10.1086/285610. Publisher: The University of Chicago Press.

27. Weller, C. & Wu, M. A generation-time effect on the rate of molecular evolution in bacteria. Evolution 69, 643–652 (2015).

28. Shoemaker, W. R. et al. Microbial population dynamics and evolutionary outcomes under extreme energy limitation. Proceedings of the National Academy of Sciences 118 (2021). URL https://www.pnas.org/content/118/33/e2101691118. Publisher: National Academy of Sciences Section: Biological Sciences.

29. Sachidanandham, R. & Yew-Hoong Gin, K. A dormancy state in nonspore-forming bacteria. Applied Microbiology and Biotechnology 81, 927–941 (2009).

30. Siebring, J. et al. Repeated triggering of sporulation in Bacillus subtilis selects against a protein that affects the timing of cell division. The ISME Journal 8, 77–87 (2014). URL https://www.nature.com/articles/ismej2013128. Bandiera_abtest: a Cg_type: Nature Research Journals Number: 1 Primary _atype: Research Publisher: Nature Publishing Group Subject_term: Bacterial physiology;Evolutionary genetics Subject_term_id: bacterial-physiology;evolutionary-genetics.

31. Zeigler, D. R. & Nicholson, W. L. Experimental evolution of Bacillus subtilis. Environmental Microbiology 19, 3415–3422 (2017).

32. Maughan, H., Masel, J., Birky, C. W. & Nicholson, W. L. The Roles of Mutation Accumulation and Selection in Loss of Sporulation in Experimental Populations of Bacillus subtilis. Genetics 177, 937–948 (2007). URL https://www.ncbi.nlm.nih.gov/pmc/articles/PMC2034656/.

33. Maughan, H. et al. The Population Genetics of Phenotypic Deterioration in Experimental Populations of Bacillus Subtilis. Evolution 60, 686–695 (2006). URL https://onlinelibrary.wiley.com/doi/abs/10.1111/j.0014-3820.2006.tb01148.x.

34. Maughan, H., Birky, C. W. & Nicholson, W. L. Transcriptome Divergence and the Loss of Plasticity in Bacillus subtilis after 6,000 Generations of Evolution under Relaxed Selection for Sporulation. Journal of Bacteriology 191, 428–433 (2009). URL https://jb.asm.org/content/191/1/428.

35. Shoemaker, W. R., Polezhaeva, E., Givens, K. B. & Lennon, J. T. Molecular Evolutionary Dynamics of Energy Limited Microorganisms. Molecular Biology and Evolution (2021). URL https://doi.org/10.1093/molbev/msab195.

36. Mason, C. A. & Hamer, G. Cryptic growth in Klebsiella pneumoniae. Applied Microbiology and Biotechnology 25, 577–584 (1987). URL https://doi.org/10.1007/BF00252019.

37. Banks, M. K. & Bryers, J. D. Cryptic growth within a binary microbial culture. Applied Microbiology and Biotechnology 33, 596–601 (1990). URL https://doi.org/10.1007/BF00172558.

38. Bradley, J. A., Amend, J. P. & LaRowe, D. E. Necromass as a Limited Source of Energy for Microorganisms in Marine Sediments. Journal of Geophysical Research: Biogeosciences 123, 577–590 (2018). URL https://agupubs.onlinelibrary.wiley.com/doi/abs/10.1002/2017JG004186.

39. Martin, M. Cutadapt removes adapter sequences from high-throughput sequencing reads. EMBnet.journal 17, 10–12 (2011). URL http://journal.embnet.org/index.php/embnetjournal/article/view/200.

40. Good, B. H., McDonald, M. J., Barrick, J. E., Lenski, R. E. & Desai, M. M. The dynamics of molecular evolution over 60,000 generations. Nature 551, 45–50 (2017). URL https://www.nature.com/articles/nature24287.

41. Deatherage, D. E. & Barrick, J. E. Identification of mutations in laboratory-evolved microbes from next-generation sequencing data using breseq. Methods in Molecular Biology (Clifton, N.J.) 1151, 165–188 (2014).

42. Transtrum, M. K. & Qiu, P. Bridging Mechanistic and Phenomenological Models of Complex Biological Systems. PLoS Computational Biology 12 (2016). URL https://www.ncbi.nlm.nih.gov/pmc/articles/PMC4871498/.

43. Seabold, S. & Perktold, J. statsmodels: Econometric and statistical modeling with python. In 9th Python in Science Conference (2010).

44. McDonald, M. J., Rice, D. P. & Desai, M. M. Sex Speeds Adaptation by Altering the Dynamics of Molecular Evolution. Nature 531, 233–236 (2016). URL https://www.ncbi.nlm.nih.gov/pmc/articles/PMC4855304/.

45. Grimmett, G. & Stirzaker, D. Probability and random processes (Oxford University Press, Oxford; New York, 2001), 3rd ed edn.

46. Patefield, W. M. Algorithm as 159: An efficient method of generating random r ×c tables with given row and column totals. Journal of the Royal Statistical Society. Series C (Applied Statistics) 30, 91–97 (1981). URL https://www.jstor.org/stable/2346669.

47. Baak, M., Koopman, R., Snoek, H. & Klous, S. A new correlation coefficient between categorical, ordinal and interval variables with pearson characteristics (2019). 1811.11440.

48. Arrieta-Ortiz, M. L. et al. An experimentally supported model of the *Bacillus subtilis* global transcriptional regulatory network. Molecular Systems Biology 11, 839 (2015). URL https://onlinelibrary.wiley.com/doi/10.15252/msb.20156236.

49. Shoemaker, W. R. & Lennon, J. T. Quantifying parallel evolution. bioRxiv 2020.05.13.070953 (2020). URL https://www.biorxiv.org/content/10.1101/2020.05.13.070953v1.

50. Good, B. H., Rouzine, I. M., Balick, D. J., Hallatschek, O. & Desai, M. M. Distribution of fixed beneficial mutations and the rate of adaptation in asexual populations. Proceedings of the National Academy of Sciences 109, 4950–4955 (2012). URL https://www.pnas.org/content/109/13/4950. Publisher: National Academy of Sciences Section: Biological Sciences.

51. Blath, J., Buzzoni, E., Casanova, A. G. & Wilke-Berenguer, M. Separation of time-scales for the seed bank diffusion and its jump-diffusion limit (2019). 1903.11795.

52. Blath, J., Buzzoni, E., Koskela, J. & Berenguer, M. W. Statistical tools for seed bank detection. arXiv (2019). URL https://arxiv.org/abs/1907.13549v3.

53. Sung, W. et al. Asymmetric Context-Dependent Mutation Patterns Revealed through Mutation-Accumulation Experiments. Molecular Biology and Evolution 32, 1672–1683 (2015). URL https://www.ncbi.nlm.nih.gov/pmc/articles/PMC4476155/.

54. Good, B. H. & Desai, M. M. The impact of macroscopic epistasis on long-term evolutionary dynamics. Genetics 199, 177–190 (2015). URL http://www.genetics.org/lookup/doi/10.1534/genetics.114.172460.

55. Ewens, W. J. Mathematical Population Genetics 1: Theoretical Introduction. Interdisciplinary Applied Mathematics, Mathematical Population Genetics (Springer-Verlag, New York, 2004), 2 edn. URL http://www.springer.com/gp/book/9780387201917.

56. Cvijović, I., Good, B. H. & Desai, M. M. The Effect of Strong Purifying Selection on Genetic Diversity. Genetics 209, 1235–1278 (2018). URL https://doi.org/10.1534/genetics.118.301058.

57. Fisher, D. S. Course 11 Evolutionary dynamics. In Les Houches, vol. 85, 395–446 (Elsevier, 2007). URL https://linkinghub.elsevier.com/retrieve/pii/S0924809907800187.

58. Good, B. H. Linkage disequilibrium between rare mutations. bioRxiv 2020.12.10.420042 (2020). URL https://www.biorxiv.org/content/10.1101/2020.12.10.420042v1. Publisher: Cold Spring Harbor Laboratory Section: New Results.

59. Schultz, D., Wolynes, P. G., Jacob, E. B. & Onuchic, J. N. Deciding fate in adverse times: Sporulation and competence in Bacillus subtilis. Proceedings of the National Academy of Sciences 106, 21027–21034 (2009). URL https://www.pnas.org/content/106/50/21027. Publisher: National Academy of Sciences Section: Physical Sciences.

60. Hahn, J., Roggiani, M. & Dubnau, D. The major role of Spo0A in genetic competence is to downregulate abrB, an essential competence gene. Journal of Bacteriology 177, 3601–3605 (1995). URL https://www.ncbi.nlm.nih.gov/pmc/articles/PMC177070/.

61. Albano, M., Hahn, J. & Dubnau, D. Expression of competence genes in Bacillus subtilis. Journal of Bacteriology 169, 3110–3117 (1987).

62. Dubnau, D. Genetic competence in Bacillus subtilis. Microbiological Reviews 55, 395–424 (1991). URL https://www.ncbi.nlm.nih.gov/pmc/articles/PMC372826/.

63. Hoch, J. A. Genetic analysis of pleiotropic negative sporulation mutants in Bacillus subtilis. Journal of Bacteriology 105, 896–901 (1971).

64. Rozen, D. E., Philippe, N., Arjan de Visser, J., Lenski, R. E. & Schneider, D. Death and cannibalism in a seasonal environment facilitate bacterial coexistence. Ecology Letters 12, 34–44 (2009). URL http://doi.wiley.com/10.1111/j.1461-0248.2008.01257.x.

65. Bradshaw, W. E., Armbruster, P. A. & Holzapfel, C. M. Fitness consequences of hibernal diapause in the pitcher-plant mosquito, wyeomyia smithii. Ecology 79, 1458–1462 (1998). URL https://www.jstor.org/stable/176758.

66. Cáceres, C. E. & Tessier, A. J. To sink or swim: Variable diapause strategies among daphnia species. Limnology and Oceanography 49, 1333–1340 (2004).

67. Dubnau, E. J.et al. A protein complex supports the production of Spo0A-P and plays additional roles for biofilms and the K-state in Bacillus subtilis. Molecular Microbiology 101, 606–624 (2016).

68. Pedrido, M. E. et al. Spo0A links de novo fatty acid synthesis to sporulation and biofilm development in Bacillus subtilis. Molecular Microbiology 87, 348–367 (2013).

69. Hamon, M. A. & Lazazzera, B. A. The sporulation transcription factor Spo0A is required for biofilm development in Bacillus subtilis. Molecular Microbiology 42, 1199–1209 (2001). URL https://onlinelibrary.wiley.com/doi/abs/10.1046/j.1365-2958.2001.02709.x. _eprint: https://onlinelibrary.wiley.com/doi/pdf/10.1046/j.1365-2958.2001.02709.x.

70. Takada, H. et al. Cell motility and biofilm formation in Bacillus subtilis are affected by the ribosomal proteins, S11 and S21. Bioscience, Biotechnology, and Biochemistry 78, 898–907 (2014). URL https://doi.org/10.1080/09168451.2014.915729.

71. Verhamme, D. T., Murray, E. J. & Stanley-Wall, N. R. DegU and Spo0A Jointly Control Transcription of Two Loci Required for Complex Colony Development by Bacillus subtilis. Journal of Bacteriology 191, 100–108 (2009). URL https://journals.asm.org/doi/full/10.1128/JB.01236-08. Publisher: American Society for Microbiology.

72. Mirouze, N., Desai, Y., Raj, A. & Dubnau, D. Spo0A P Imposes a Temporal Gate for the Bimodal Expression of Competence in Bacillus subtilis. PLOS Genetics 8, e1002586 (2012). URL https://journals.plos.org/plosgenetics/article?id=10.1371/journal.pgen.1002586. Publisher: Public Library of Science.

73. Wetmore, K. M. et al. Rapid Quantification of Mutant Fitness in Diverse Bacteria by Sequencing Randomly Bar-Coded Transposons. mBio 6 (2015). URL https://mbio.asm.org/content/6/3/e00306-15. Publisher: American Society for Microbiology Section: Research Article.

74. Wright, E. S. & Vetsigian, K. H. Stochastic exits from dormancy give rise to heavy-tailed distributions of descendants in bacterial populations. Molecular Ecology 28, 3915–3928 (2019). URL https://onlinelibrary.wiley.com/doi/abs/10.1111/mec.15200. _eprint: https://onlinelibrary.wiley.com/doi/pdf/10.1111/mec.15200.

75. Kinsler, G., Geiler-Samerotte, K. & Petrov, D. A. Fitness variation across subtle environmental perturbations reveals local modularity and global pleiotropy of adaptation. eLife 9, e61271 (2020). URL https://doi.org/10.7554/eLife.61271. Publisher: eLife Sciences Publications, Ltd.

76. Reddy, G. & Desai, M. M. Global epistasis emerges from a generic model of a complex trait. eLife 10, e64740 (2021). URL https://doi.org/10.7554/eLife.64740. Publisher: eLife Sciences Publications, Ltd.

