## Supplemental Information for "Seed banks alter the molecular evolutionary dynamics of *Bacillus subtilis*"

January 14, 2022

### 1 *B. subtilis* ∆spo0A mutant construction

#### 1.1 Justification

While the ability to regulate metabolic activity can be viewed as a quantitative trait, as a quantity, it is difficult to experimentally manipulate. Instead, we chose to use *B. subtilis*, an organism that forms a physiological structure in response to resource depletion. This adaptation can be manipulated as a binary variable, which we accomplished by deleting *spo0A*, the master regulatory gene for sporulation pathways in *B. subtilis*.

#### 1.2 Plasmid construction

Plasmid pMiniMAD ∆*spo0A* was constructed by PCR amplifying the 1,000 bp genomic regions upstream and downstream of the *spo0A* gene using custom primers (Table S1). Additional nucleotides at the end of the upstream and downstream primers were added for complementation with the plasmid *smaI* restriction site. The two PCR amplified dsDNA fragments (*Spo0A-U* and *Spo0A-D*) were purified using a QiaGen QIAquick PCR Purification Kit.

#### 1.3 Gibson assembly

The two PCR amplified dsDNA fragments (*Spo0A-U* and *Spo0A-D*) were ligated into a pMiniMAD plasmid using the one-step isothermal DNA assembly method (Gibson Assembly). Master mix containing 15 $\mu$L of buffer-enzymes mix, 3 $\mu$L of pMiniMAD plasmid solution, 1 $\mu$L of *Spo0A-U* and 1 $\mu$L *Spo0A-D* DNA fragments. The reaction was carried out in the thermocycler at 50°C for 1 hour. The presence of the product was confirmed using gel electrophoresis. The product was purified by drop dialysis against molecular grade water for 30 min.

#### 1.4 Transformation

The ligated plasmid was transformed into *E. coli* DH5$\alpha$ as follows: 50 $\mu$L cell suspension by electroporation, diluted with 1 mL of of LB media, incubated in shaker at 37°C for 1 hour. Positive transformants were selected on LB-agar containing 100 $\mu$g/mL ampicillin overnight at 37°C. Plasmid DNA was purified from 1 mL of culture using a QiaGen MiniPrep Kit. To confirm that new fragments were ligated to pMiniMad plasmid, a solution of plasmid was treated with restriction enzymes (5 $\mu$L of plasmid solution, 2 $\mu$L of CutSmart buffer, 0.5 $\mu$L EcoRI-HF (enzymes), 12.5 $\mu$L water) at 37°C for 1 hour and analyzed using gel electrophoresis.

#### 1.5 *E. coli* TG1 transformation

Purified plasmid was transformed into *E. coli* TG1 following the same procedure (2$\mu$L of plasmid solution, 50 $\mu$L of *E. coli* suspension). Positive transformants were selected on LB-agar containing 100 $\mu$g/mL ampicillin (37°C, overnight).

Plasmid DNA was purified from 1 ml of overnight culture (3 ml of LB+100 $\mu$g/mL ampicillin) using QiaGen MiniPrep Kit and stored at -20°C.

##### 1.6 *B. subtilis* NCIB 3610 transformation

A *B. subtilis* colony was inoculated into 2 mL of MC medium and grown at 37°C for 4.5 hours. (MC media: 1.8 mL of water, 0.2 mL of 10xMC, 20 $\mu$l of 0.3M $MgSO_4$). A 4 $\mu$L aliquot of the purified plasmid was added to 400 $\mu$L of *B. subtilis* culture and incubated in shaker at 37°C for 2 hrs. and plated on LB+mls agar. Positive transformants were identified by selecting for resistance to macrolide-lincosamide-streptogramin (MLS) on LB+MLS agar and colonies were streaked on the LB+MLS agar again. Mutants were identified by loss of antibiotic resistance. Colony PCR was run using the Spo0AUF ITA and Spo0ADR ITA primers for 3 colonies and mutants were identified using a gel to confirm that the amplified fragment was ∼2kb. Loss of antibiotic resistance was confirmed by growing the same colony on LB+MLS agar.

#### 2 Two locus evolution with a seed bank

##### 2.1 Theoretical motivation

We performed additional simulations to determine the degree that a purported lower recombination rate in $\Delta$*spo0A* could explain the Kolmogorov-Smirnov distance of distributions of mutation trajectory measures between populations that can and cannot form seed banks. To provide a framework grounded in first principles from theoretical population genetics, we constructed a mathematical model of the evolution of two loci in the presence of a seed bank. Our simulations, and the theory that grounds them, were motivated by recent developments on the dynamics of linked mutations [1]. To start, we consider the frequencies of four possible haplotypes in a population harboring two loci: $f_{11}$, $f_{10}$, $f_{01}$, and $f_{00}$. In the diffusion limit, the dynamics of all four haplotype frequencies can be described as a coupled system of nonlinear Langevin equations. In addition, an individual of a given haplotype can be active ($A$) or dormant ($D$) at a given generation, meaning that the full system contains eight equations (four genotypes * two demographic states). Active individuals are subject to selection, mutation, recombination, random genetic drift, resuscitation, and dormancy, whereas dormant individuals are only subject to resuscitation and dormancy.

$$\frac{\partial f_{00}^A}{\partial t} = [RC_{\mathrm{LD}}(t) + \frac{\xi_{00}(t)}{\sqrt{A}} \tag{S1a}$$

$$+ \mu(f_{10}^A + f_{01}^A) - 2\mu f_{00}^A - cf_{00}^A + cKf_{00}^D \tag{S1b}$$

$$\frac{\partial f_{10}^A}{\partial t} = s_{10} f_{10}^A \left(\frac{A}{A+D} - f_{10}^A\right) + RC_{\mathrm{LD}}(t) + \frac{\xi_{10}(t)}{\sqrt{A}} \tag{S1c}$$

$$+ \mu(f_{00}^A + f_{11}^A) - 2\mu f_{10}^A - cf_{10}^A + cKf_{10}^D \tag{S1d}$$

$$\frac{\partial f_{01}^A}{\partial t} = s_{01} f_{01}^A \left(\frac{A}{A+D} - f_{01}^A\right) + RC_{\mathrm{LD}}(t) + \frac{\xi_{01}(t)}{\sqrt{A}} \tag{S1e}$$

$$+ \mu(f_{00}^A + f_{11}^A) - 2\mu f_{01}^A - cf_{01}^A + cKf_{01}^D \tag{S1f}$$

$$\frac{\partial f_{11}^A}{\partial t} = s_{11} f_{11}^A \left(\frac{A}{A+D} - f_{11}^A\right) - RC_{\mathrm{LD}}(t) + \frac{\xi_{11}(t)}{\sqrt{A}} \tag{S1g}$$

$$+ \mu(f_{10}^A + f_{01}^A) - 2\mu f_{11}^A - cf_{11}^A + cKf_{11}^D \tag{S1h}$$

$$\frac{\partial f_{00}^D}{\partial t} = cf_{00}^A - cKf_{00}^D \tag{S1i}$$

$$\frac{\partial f_{10}^D}{\partial t} = cf_{10}^A - cKf_{10}^D \tag{S1j}$$

$$\frac{\partial f_{01}^D}{\partial t} = cf_{01}^A - cKf_{01}^D \tag{S1k}$$

$$\frac{\partial f_{11}^D}{\partial t} = cf_{11}^A - cKf_{11}^D \tag{S1l}$$

where $s_{10}$, $s_{01}$, and $s_{11}$ are the selection coefficients for each genotype, with $s_{00} = 0$. The standard coefficient of linkage disequilibrium is defined as $C_{\mathrm{LD}}(t) \equiv f_{11}^A f_{00}^A - f_{10}^A f_{01}^A$. Because recombination only occurs between active individuals, the frequencies of dormant haplotypes do not alter the degree of disequilibrium between haplotypes. Likewise, because dormant individuals do not reproduce, the maximum obtainable frequency of a beneficial mutation due to selection alone is equal to the fraction of reproducing hosts in the population ($\frac{A}{A+D}$) rather than one (i.e., all individuals are able to reproduce). The terms $\xi_i(t)$ are Brownian noise terms that capture the stochastic fluctuations due to random genetic drift [2].

##### 2.2 Simulations

We performed forward-time simulations based on the dynamics in Eq.S1l. Following a previous two-locus simulation approach [1], we used a Wright-Fisher sampling scheme. We examined the $A\mu \ll 1$ limit, allowing us to drop the mutation terms in our system of equations. To improve computational efficiency and to examine the impact of recombination when it is most relevant, we performed our simulations when both mutations were simultaneously segregating. This was done by drawing an initial single-locus haplotype frequency from the site frequency spectrum of a population with a seed bank [3]

$$p(f|s,D,c) \propto \frac{(D/c)^2}{f(1-f)} \frac{1-\exp[-2Ns(D/c)(1-f)]}{1-\exp[-2Ns(D/c)]} \tag{S2}$$

We then introduced a second mutation to an active individual harboring the mutant or the ancestor with with probability $f$ or $1-f$, respectively. A single simulation proceeded until one of the two mutations went extinct. We recorded our measures of mutation trajectories in Eq.4. every $\Delta t = \Delta \tau = 75$ generations, a timescale roughly equal to the number of generations that accrued in the 10-day WT populations between sequenced timepoints. We performed 1,000 iterations of the simulation for each parameter combination. We performed bootstrap sampling of the simulated distributions to determine whether the Kolmogorov-Smirnov distance between dormancy-capable and incapable populations for a given mutation trajectory statistic increased with the population scaled recombination rate $A \cdot R$.

#### References


[1] Benjamin H. Good. "Linkage disequilibrium between rare mutations". en. In: *bioRxiv* (Dec. 2020). Publisher: Cold Spring Harbor Laboratory Section: New Results, p. 2020.12.10.420042. DOI: 10.1101/2020.12.10.420042. URL: https://www.biorxiv.org/content/10.1101/2020.12.10.420042v1 (visited on 08/02/2021).

[2] Benjamin H. Good and Michael M. Desai. "Fluctuations in fitness distributions and the effects of weak linked selection on sequence evolution". en. In: *Theoretical Population Biology* 85 (May 2013), pp. 86–102. ISSN: 0040-5809. DOI: 10.1016/j.tpb.2013.01.005. URL: https://www.sciencedirect.com/science/article/pii/S0040580913000063 (visited on 12/07/2021).

[3] Bendix Koopmann et al. "Fisher-Wright model with deterministic seed bank and selection". eng. In: *Theoretical Population Biology* 114 (2017), pp. 29–39. ISSN: 1096-0325. DOI: 10.1016/j.tpb.2016.11.005.

[4] William R Shoemaker et al. "Molecular Evolutionary Dynamics of Energy Limited Microorganisms". In: *Molecular Biology and Evolution* msab195 (July 2021). ISSN: 0737-4038. DOI: 10.1093/molbev/msab195. URL: https://doi.org/10.1093/molbev/msab195 (visited on 08/24/2021).

#### 3 Tables

| Primer | Sequence |
|---|---|
| spo0AUF ITA | 5'-GTCGACTCTAGAGGATCCCCACTGATCAAACGTGATAAACAG-3' |
| spo0AUR ITA | 5'-CAATGAATTCACTGTTGGTAGGCAGGCTTACCAGCTCTCGATT-3' |
| spo0ADF ITA | 5'-AATCGAGAGCTGGTAAGCCTGCCTACCAACAGTGAATTCATTG-3' |
| spo0ADR ITA | 5'-GTGAATTCGAGCTCGGTACCCCGGAAGAACCTGAGACACC-3' |

Table S1: PCR primers described in the methods section of the main manuscript.

| Transfer regime | Strain | $[\log_{10} M]_{\max} \pm$ S.E. | $t_{1/2} \pm$ S.E. [gens.] |
|---|---|---|---|
| 1-day | WT | $2.14 \pm 0.190$ | $502.1 \pm 150.4$ |
| | $\Delta$*spo0A* | $1.73 \pm 0.186$ | $511.0 \pm 175.1$ |
| 10-days | WT | $4.67 \pm 3.97$ | $1,865 \pm 2,171$ |
| | $\Delta$*spo0A* | $1.70 \pm 0.361$ | $64.33 \pm 47.26$ |
| 100-days | WT | $7.87 \pm 22.0$ | $422.1 \pm 1,358$ |
| | $\Delta$*spo0A* | $1.08 \pm 0.661$ | $27.32 \pm 51.13$ |

Table S2: Parameters estimated by fitting Eq.3 to empirical data via numerical optimization. Standard errors of the estimates (S.E.) were obtained by calculating the square root of the variance-covariance matrix of parameter estimates.

| Transfer regime | Strain | $t$, *spo0A* vs. non-*spo0A* | $P$ |
|---|---|---|---|
| 1-day | WT | $-0.842$ | 0.447 |
| | $\Delta$*spo0A* | $-1.89$ | 0.131 |
| 10-days | WT | 1.04 | 0.358 |
| | $\Delta$*spo0A* | 0.513 | 0.635 |
| 100-days | WT | 2.40 | 0.0744 |
| | $\Delta$*spo0A* | $-1.40$ | 0.235 |

Table S3: The ratio of the fraction of nonsynonymous to synonymous polymorphisms was consistently less than one across strains and treatments (Fig. S4), a pattern indicative of widespread purifying selection. To determine whether the removal of *spo0A* in the $\Delta$*spo0A* mutant induced relaxed selection on the set of genes regulated by *spo0A* (i.e., the regulon), $pN/pS$ was estimated for the set of genes in the regulon and the set of genes outside the regulon for each replicate population. To test whether $pN/pS$ was higher outside the regulon, paired one-way $t$-tests were performed for each strain-transfer combination. We found no evidence that $pN/pS$ was higher in the regulon within a given strain-transfer combination.

| Transfer regime | Strain | $\Delta\ell$ | $P$ |
|---|---|---|---|
| 1-day | WT | 2.34 | $< 10^{-5}$ |
| | $\Delta$*spo0A* | 2.15 | $< 10^{-5}$ |
| 10-days | WT | 3.23 | $< 10^{-5}$ |
| | $\Delta$*spo0A* | 1.31 | $< 10^{-5}$ |
| 100-days | WT | 2.59 | $< 10^{-5}$ |
| | $\Delta$*spo0A* | 2.38 | $< 10^{-5}$ |

Table S4: Likelihood of enrichment of nonsynonymous mutations across genes ($\Delta\ell$) and significance for all strain-transfer regime combinations.

| Transfer regime | Strain | $n_{\mathrm{regulon}}$ | $P$ |
|---|---|---|---|
| 1-day | WT | 113 | 0.0466 |
| | $\Delta$*spo0A* | 33 | 0.00930 |
| 10-days | WT | 37 | 0.996 |
| | $\Delta$*spo0A* | 58 | 0.199 |
| 100-days | WT | 45 | 0.925 |
| | $\Delta$*spo0A* | 26 | 0.0648 |

Table S5: The number of nonsynonymous mutations acquired within the *spo0A* regulon for a given strain-transfer regime ($n_{\mathrm{regulon}}$). $P$-values were obtained using the binomial distribution where the probability of a mutation landing in the regulon is equal to the proportion of nonsynonymous sites that the regulon contains.

| Transfer regime | Enriched strain | $\|\Delta p_{\text{regulon}}\|$ | $P$ |
|---|---|---|---|
| 1-day | $\Delta$*spo0A* | 0.0110 | 0.160 |
| 10-days | $\Delta$*spo0A* | 0.0160 | 0.007 |
| 100-days | $\Delta$*spo0A* | 0.0170 | 0.038 |

Table S6: The absolute difference in the fraction of nonsynonymous mutations acquired in genes that are in the *spo0A* regulon between WT vs. $\Delta$*spo0A* strains for all transfer regimes ($|\Delta p_{\text{regulon}}|$). $P$-values were obtained by binomial sampling where the probability of a mutation landing on a nucleotide in the regulon is equal to the fraction of nonsynonymous bases that are in genes within the regulon.

| Transfer regime | Enriched strain | $\lvert\Delta p_{\mathrm{spore}}\rvert$ | $P$ |
|---|---|---|---|
| 1-day | WT | 0.0313 | $10^{-5}$ |
| 10-days | WT | 0.104 | $10^{-5}$ |
| 100-days | WT | 0.0623 | $10^{-5}$ |

Table S7: The absolute difference in the fraction of mutations acquired in genes that encode for endospore formation between WT and Δ*spo0A* strains ($\lvert\Delta p_{\mathrm{spore}}\rvert$). $P$-values were obtained by binomial sampling where the probability of a mutation landing on an endospore encoding nucleotide is the fraction of nonsynonymous bases that are in genes that encode for endospore formation.

#### 4 Figures

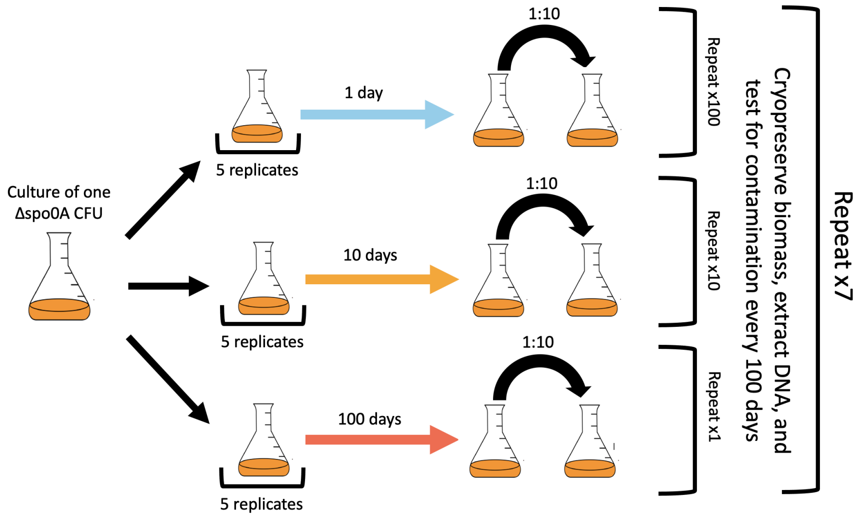


Figure S1: A conceptual diagram illustrating the experimental evolution protocol. A single colony isolate of Δ*spo0A* was grown overnight and split into 15 flasks. Three sets of five flasks were transferred every 1, 10, or 100 days for 700 days, corresponding to 700, 70, and 7 transfers, respectively. All sample collection took place every 100 days for all replicate populations. Figure modified from a prior publication [4].

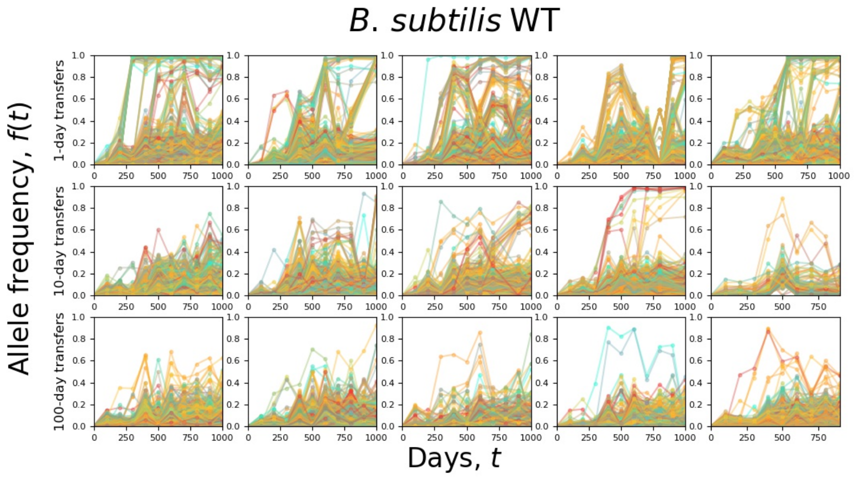


Figure S2: Allele frequency trajectories of all *Bacillus subtilis* WT replicate populations.

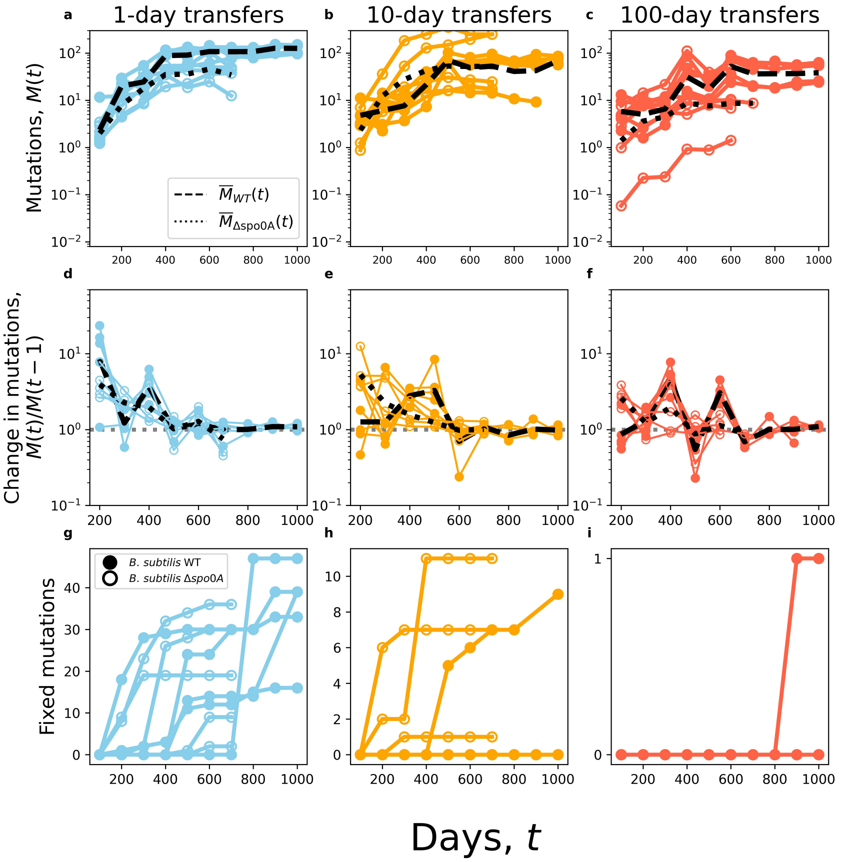


Figure S3: Molecular evolutionary dynamics of *B subtilis* WT and $\Delta spo0A$. **a-c)** Cumulative mutation ($M(t)$) trajectories for both strains over time. The dashed black line is the mean for wt populations and the dotted black line is the mean for $\Delta spo0A$. Closed circles represent wt populations and open circles represent $\Delta spo0A$ populations. **d-f)** The change in $M(t)$ between timepoints across treatments Generally there is a rapid decrease in the rate of $M(t)$, followed by it settling to an equilibrium value by day 500. **g-i)** The cumulative number of fixed mutations over time for all WT and $\Delta spo0A$ populations.

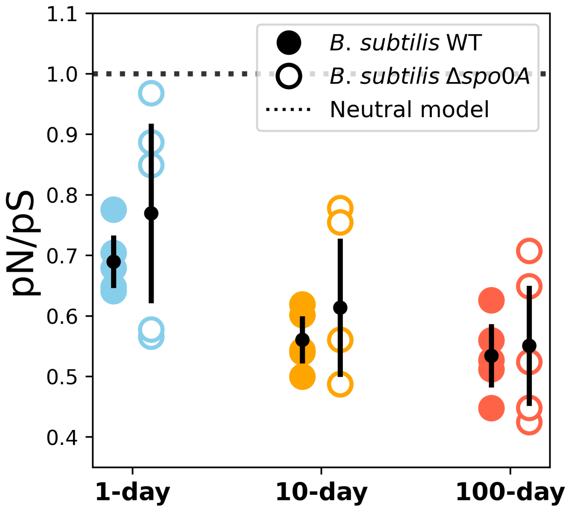


Figure S4: While the ratio of nonsynonymous to synonymous mutations ($pN/pS$) is consistently less than one across treatments (black dashed line), it does not vary between *B. subtilis* WT and $\Delta$*spo0A* populations.

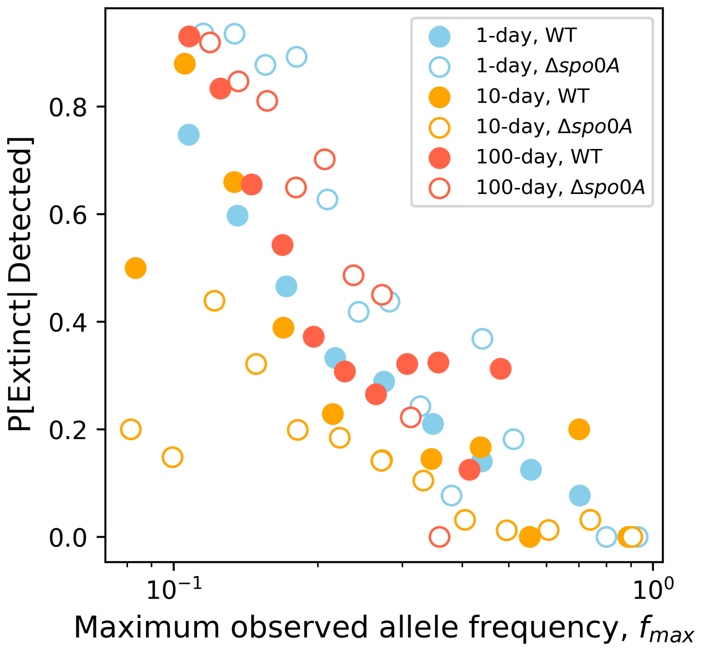


Figure S5: Given the absence of fixation events, $f_{max}$ is arguably a sufficient proxy. To justify this argument, we examined the empirical relationship between $f_{max}$ and the complement of the probability of fixation: extinction. If $f_{max}$ reflects the probability of fixation of a mutation than we should observe an inverse relationship between $f_{max}$ and the probability of extinction. We find that such a relationship exists for all strain-transfer regime combinations, allowing us to proceed with our analysis of the distribution of $f_{max}$.

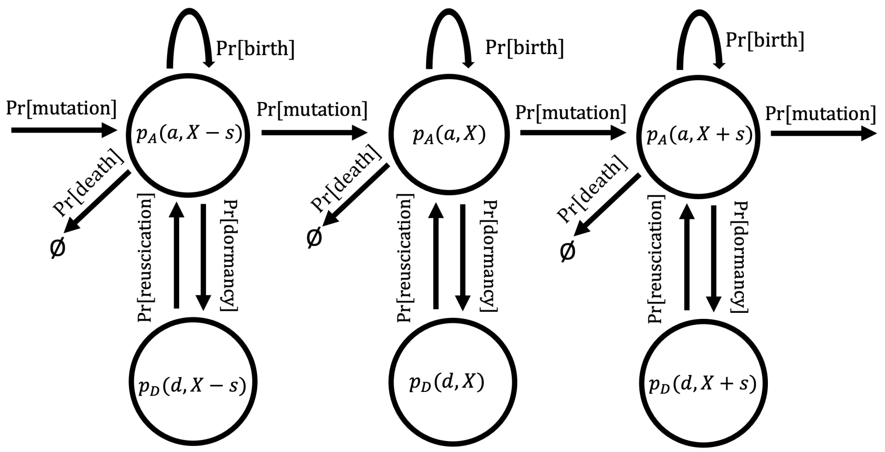


Figure S6: A conceptual diagram visualizing the transition probabilities described in Eq.14. The dependence on time ($t$) has been removed from the probability terms for the purpose of visualization.

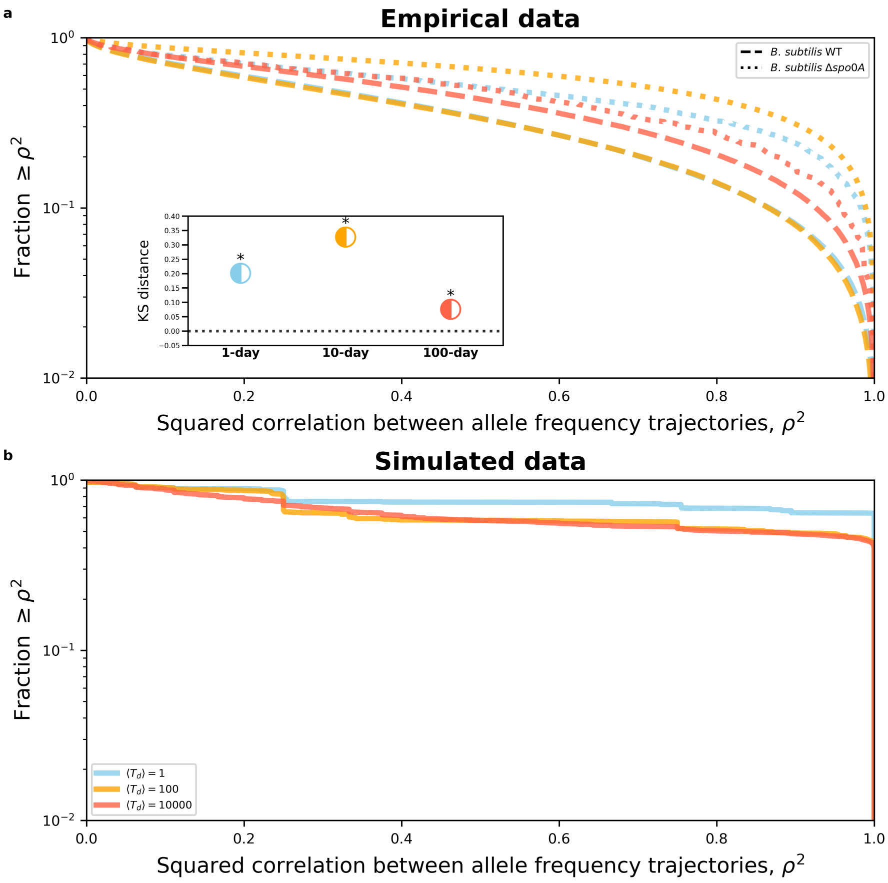


Figure S7: A decrease in the amount of recombination in a population corresponds to greater statistical dependence between mutation trajectories. This dependence can be estimated as the pairwise squared correlation of frequencies between a given pair of mutation trajectories ($\rho^2$), where a higher value of $\rho^2$ indicates a lower degree of recombination. **a)** The $\Delta$*spo0A* strain typically had a higher value of $\rho^2$ across transfer regimes, suggesting a lower degree of recombination. **b)** To confirm that this observation was not driven by the presence of absence of a seed bank, we estimated $\rho^2$ from allele frequency trajectories that were obtained by simulating Eq.14.

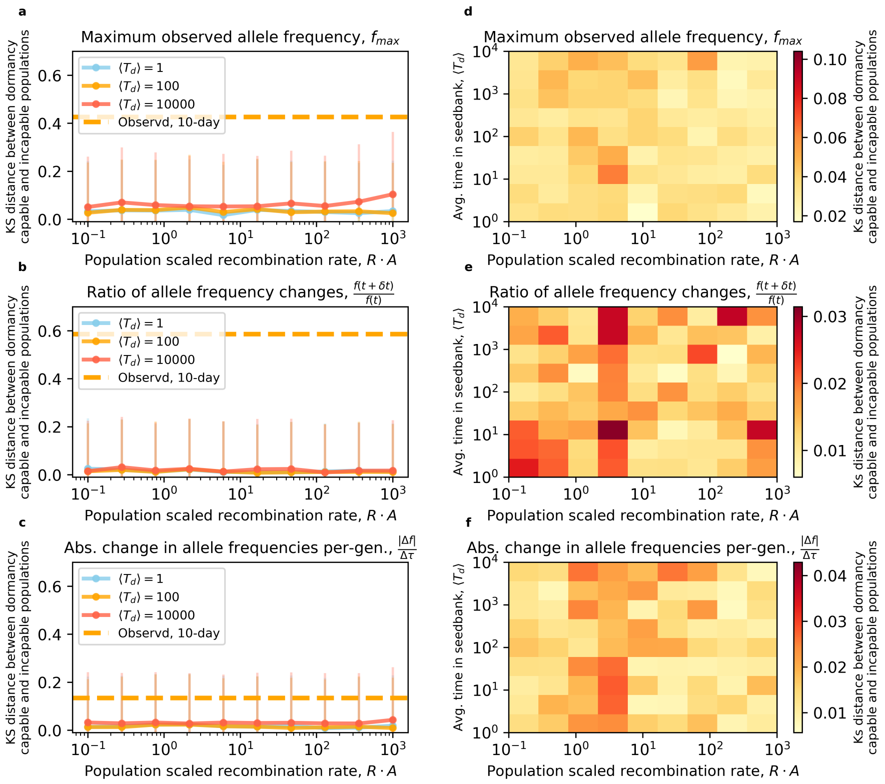


Figure S8: Two-locus simulations were performed using Eq.S1 to determine the extent that a supposed difference in the rate of recombination between the WT and $\Delta$*spo0A* could explain the distributions of mutation trajectory statistics in Fig. 3 **a-c)**. The Kolmogorov-Smirnov distance between dormancy capable and incapable populations for a given mutation trajectory statistic does not exhibit visible change over five orders of magnitude of variation in the population scaled recombination rate $RA$. This consistent lack of trend means that a competence-induced lack of recombination in $\Delta$*spo0A* cannot explain the higher KS distance that is consistently observed between WT and $\Delta$*spo0A* in the 10-day transfer regime for all mutation trajectory statistics. **d-f)** A fine-grained examination of KS distances across a range of values of $RA$ and average time in a dormant state $\langle T_d \rangle$ further supports this conclusion, as the KS distance values and degree of dispersion remains low across parameter regimes. Parameter values of $A = 10^6$, $s_{01} = s_{10} = 0.01$, and $s_{11} + s_{01} + s_{10}$ were chosen for this simulation.

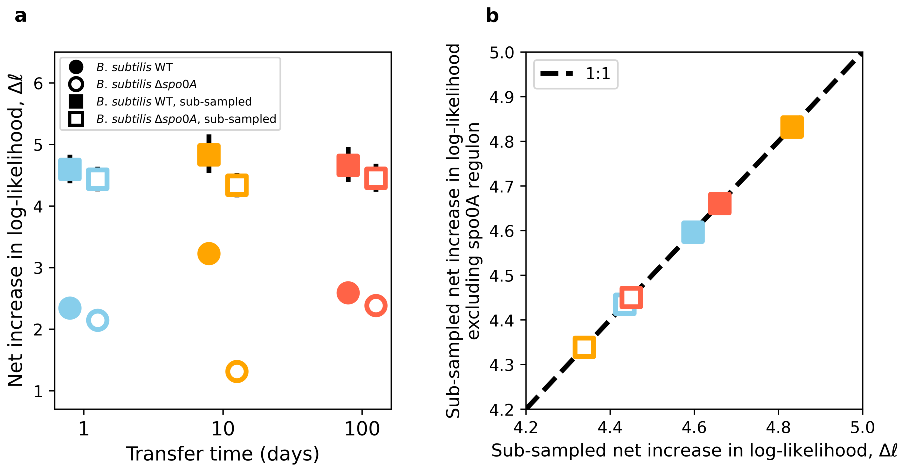


Figure S9: **a)** The genome-wide log-likelihood that certain genes were enriched for nonsynonymous mutations for *B. subtilis* WT and $\Delta$*spo0A*. Log-likelihood values for all mutations are shows as circles, where log-likelihood values for 10,000 iterations of sub-sampling for 50 mutations are shown as squares. Black bars represent 95% CIs. **b)** To determine whether the *spo0A* regulon impacted the degree of parallelism in certain strain-transfer regime combinations, we repeated the same sub-sampling likelihood analysis for the set of genes that are not part of the regulon.

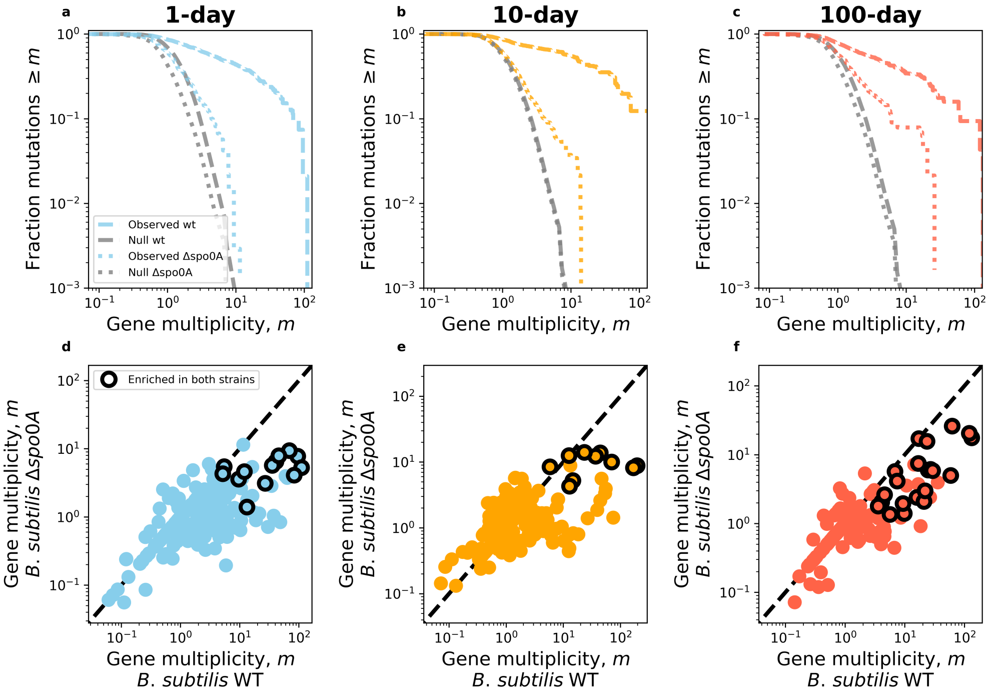


Figure S10: Parallelism analyses for *B. subtilis* WT vs. $\Delta$*spo0A* **a-c)**) The survival curves for multiplicity decays at a slower rate than the null across treatments and while $\Delta$*spo0A* is closer to the null expectation than the WT, it is significantly different from the null expectation **d-f)** The few genes that are significantly enriched for synonymous mutations in both strains within a given treatment fall near the 1:1 line.

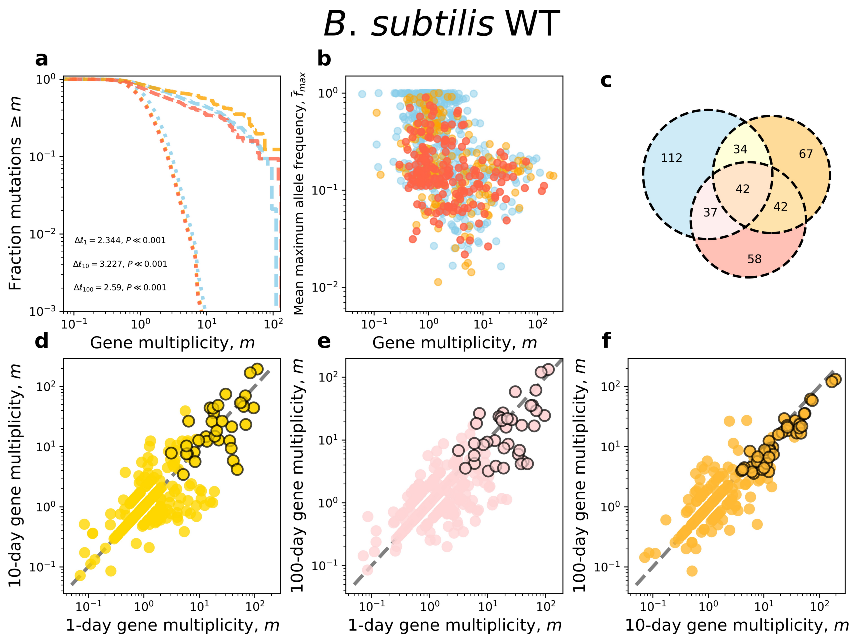


Figure S11: Parallelism analyses for *B. subtilis* WT. **a)** The survival curve for multiplicity decays at a slower rate than the null across treatments, indicating that we can reject the null hypothesis that nonsynonymous mutations are equally distributed across genes. **b)** There is no clear relationship between the multiplicity of a gene and the mean maximum frequency of mutations in the gene ($f_{max}$). **c)** Generally, significantly enriched genes tend to be enriched in more than one treatment. **d-f)** This pattern is reflected in the pairwise comparison of gene multiplicities in different treatments. This data is replotted from a prior publication for convenience [4]

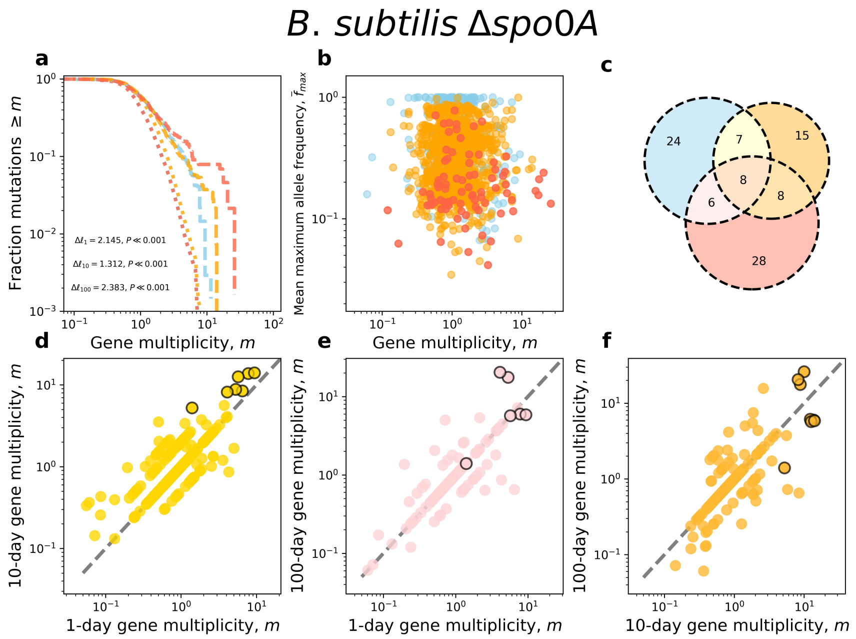


Figure S12: Parallelism analyses for *B. subtilis* Δ*spo0A* **a)** The survival curve for multiplicity decays at a slower rate than the null across treatments, indicating that we can reject the null hypothesis that nonsynonymous mutations are equally distributed across genes. **b)** There is no clear relationship between the multiplicity of a gene and the mean maximum frequency of mutations in the gene ($f_{max}$). **c)** Generally, significantly enriched genes tend to be enriched in more than one treatment. **d-f)** This pattern is reflected in the pairwise comparison of gene multiplicities in different treatments.

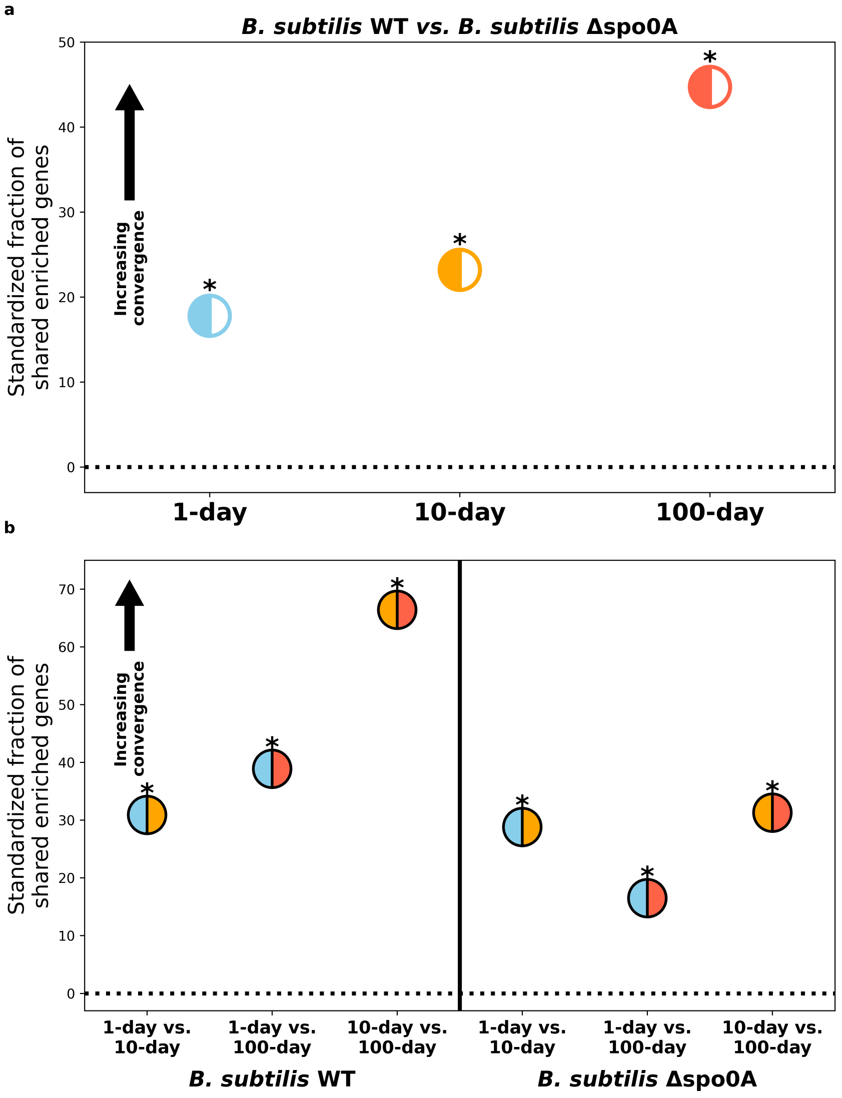


Figure S13: The degree of divergence vs. convergence as overlap in enriched genes. **a)** Similarity in the identity of enriched genes (i.e., Jaccard index) compared to a null distribution obtained via simulation indicates that convergent evolution overwhelmingly occurred between WT and Δ*spo0A* for all treatments. **b)** The same pattern was obtained for treatment comparisons within each strain.

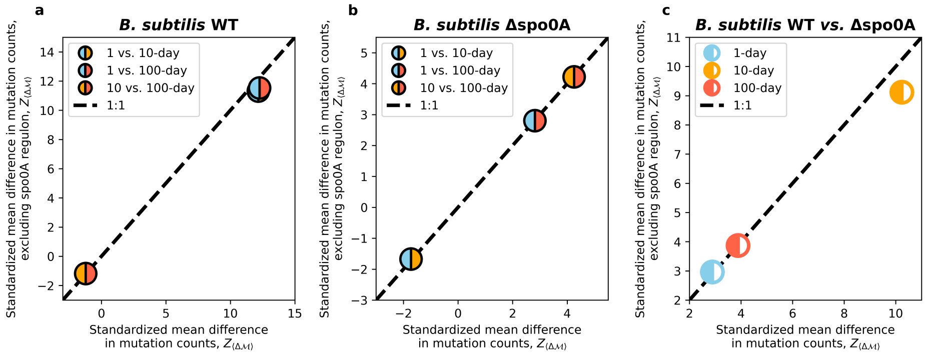


Figure S14: To determine whether the *spo0A* regulon contributed towards our convergence/divergence results, we repeated our analysis described by Eq.10 on all genes that were not part of the regulon. The effect of genes in the *spo0A* regulon was slight to nonexistent, only causing a slight deviation in our comparison of WT vs. $\Delta$*spo0A* strains in the 10-day transfer regime.

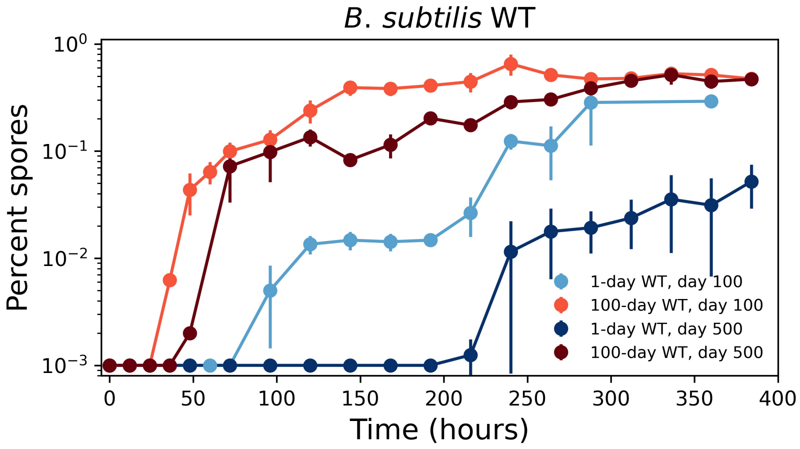


Figure S15: Endospore accumulation curves for evolved *B. subtilis* WT strains. Generally, the 100-day replicate population we chose appear to retain the general form of their survival curves over the course of the experiment (day 100 vs. 500). However, the replicate population we chose from the 1-day transfer regime form spores at a substantially slower rate at day 100. By day 500 the fraction of spores had only reached 1% over ten days. Bars represent the standard error of the mean.
